# Data-based large-scale models provide a window into the organization of cortical computations

**DOI:** 10.1101/2023.04.28.538662

**Authors:** Guozhang Chen, Franz Scherr, Wolfgang Maass

**Author notes:** Contributed equally (Guozhang Chen, Franz Scherr).

## Abstract

The neocortex of the brain is one of the most powerful computing devices that exists, but it remains largely open how its computations are organized. Since the neocortex is a 2D tapestry consisting of repeating stereotypical local cortical microcircuits, a key step for solving this problem is to understand how cortical microcircuits compute. We know by now a lot about their connectivity structure and their neuron types, but we are lacking tools for elucidating causal relations between this structure and their computational function. We present a new tool for elucidating this relation: We train large-scale models of cortical microcircuits, which integrate most current knowledge about their structure, for carrying out similar computational tasks as in the brain. We show that the trained model achieves a similar computational performance as the brain, and that it reproduces experimentally found traits of cortical computation and coding that do not appear in neural network models from AI. Furthermore, we reverse-engineer how computations are organized in the model, thereby producing specific hypotheses that can be tested in experimental neuroscience. Altogether we show that cortical microcircuits provide a distinct new neural network paradigm that is of particular interest for neuromorphic engineering because it computes with highly energy-efficient sparse activity.

**Teaser:** Reverse engineering of cortical computations

## 1. Introduction

We tackle a central problem in computational science: How does the brain compute, or more precisely how does the neocortex compute, its most prominent computational engine? A major advance came through anatomical data revealing that the mammalian neocortex is structured as a continuous 2D sheet that is stitched together from repetitions of a rather stereotypical local circuit template, commonly referred to as cortical microcircuit [Mountcastle, 1998, Douglas and Martin, 2004, Harris and Shepherd, 2015]. These cortical microcircuits are unlike neural networks that are commonly considered in AI: They consist of over 100 genetically different types of neurons that are spatially organized into 6 horizontal layers (laminae), with numerous vertical connections between specific neuron types on different layers and primarily local horizontal connections. A common hypothesis is that understanding how the structure of cortical microcircuits shapes their computations provides the key for understanding cortical computations [Mountcastle, 1998, Douglas and Martin, 2004, Harris and Shepherd, 2015].

We have by now substantial knowledge about the structure of cortical microcircuits, which results from several decades of dedicated research in several labs [Mountcastle, 1998, Thomson and Lamy, 2007, Markram et al., 2015] and subsequent further work at the Allen Institute. Furthermore, the Allen Institute has made this knowledge publicly available, in the form of a model for a representative cortical microcircuit in the primary visual cortex (area V1) of the mouse. It consists of 51,978 neurons [Billeh et al., 2020] of 111 different types. We address here the fundamental question what computational properties this experimentally found structure of the microcircuit induces.

The model of [Billeh et al., 2020] comes together with a qualitative model of the retina and LGN that transform visual stimuli into synaptic inputs to neurons in the microcircuit from V1. We will refer to this combination in the following simply as V1 model. The inclusion of the model for the retina and LGN enables us to train the V1 model to solve the same computational tasks, using the same type of natural images as used for *in vivo* experiments. We focus on two quite different tasks that have frequently been used for *in vivo* experiments: the visual-change-detection task with natural images [Garrett et al., 2020b, Siegle et al., 2021] and an evidence-accumulation-task where different visual cues have to be integrated in order to reach a decision [Morcos and Harvey, 2016a, Engelhard et al., 2019]. We trained the V1 model through stochastic gradient descent to solve these tasks, specifically leveraging a suitably modified version of BPTT (backpropagation through time). This involves pseudo-derivatives for spiking neurons as in [Bellec et al., 2018], and further modifications from [Chen et al., 2022] for the more complex generalized leaky integrate-and-fire (GLIF_3_) neuron models [Billeh et al., 2020] that had been fitted to recordings from diverse neurons from the Allen Brain Atlas [Allen Institute, 2018]. Despite the computational challenges of training such a V1 model with BPTT, advancements in software like TensorFlow [Abadi et al., 2015] and GPU hardware made this feasible.

The trained V1 model achieved for both visual processing tasks comparable performance to behavioral experiments in mice, with a very similar distribution of firing rates. Furthermore, it reproduced two characteristic traits of cortical computation that are challenging to reproduce in previously considered neural network models. First, cortical microcircuits use the timing of events to encode information, and compute with these temporal codes. More precisely, neurons display peak firing activity at specific trial times during a trial, with varying rank order based on task conditions [Driscoll et al., 2017, Koay et al., 2022, Yiling et al., 2023]. The idea to use spike timing for encoding analog values had already been proposed a long time ago [Thorpe, 1990]. However, rank order coding with single spikes is not robust to deletion or addition of single spikes. The mentioned experimental results show that the neocortex employs a less brittle rank order coding scheme: Instead of single spikes, the rank order of the times of peak firing activity of different neurons is used to encode information. We show that this type of noise-robust temporal coding can be reproduced in the data-based V1 model, but not in commonly considered types of spiking neural networks. Rank order coding is of particular interest for NMHW because it has recently been shown to provide close-to-optimal energy efficiency for neural coding [Boahen, 2022].

We found that the rank order in the V1 model contains information about the current and past visual inputs. Since this coding property has not yet been reported from *in vivo* experiments, we test this prediction by re-analyzing *in vivo* neuropixel data for the same computational task for which the model has been trained.

The second characteristic trait of cortical computations that the V1 model reproduces but which is hard to reproduce in common neural network models is their sensitivity of the network decision to the activity of very small subsets of neurons [Houweling and Brecht, 2008, Doron et al., 2014, Marshel et al., 2019, Dalgleish et al., 2020, Doron et al., 2020]. This sensitivity of the network is especially remarkable in view of the substantial trial-to-trial variability of their neural activity *in vivo*. One can see it as a special case of functional segregation of brain networks, a trait that had been proposed by [Tononi et al., 1998] to characterize, together with their functional integration capability, the complexity of brain networks. Whereas these ideas were so far primarily applied on the level of the whole brain, we show here that they are also very relevant on the scale of cortical microcircuits. We introduce quantitative measures to evaluate both rank order coding and functional segregation of cortical computations, and apply these measures to the V1 model, and also to a fairly large number of control models. The latter allows us to determine which features of the V1 model are causally linked to each characteristic trait of cortical computation.

We also analyze the flow of information in the V1 model, and use that to reverse-engineer how it computes. Its dynamics exhibit nested bifurcations, which are controlled by small segregated sets of neurons. The model even allows us to look inside these neurons, so that we can determine which features of their internal dynamics enable them to take on this pivotal role for network computations.

Altogether we show that data-based models for cortical microcircuits, trained to perform the same computational tasks as used for *in vivo* experiments, provide a fruitful new tool for understanding the organization of cortical computations. Since the energy consumption of visual processing in cortical microcircuits is by several orders of magnitude lower than that of commonly used artificial neural networks, they also provide a paradigm for a new generation of artificial neural networks with a substantially smaller energy footprint.

## 2 Results

### 2.1 Setting up a data-based V1 model for reverse engineering of cortical computations

The V1 model of [Billeh et al., 2020] represents one of the most comprehensive efforts to integrate the available experimental data on the anatomy and neurophysiology of area V1 in mouse that is currently available (Fig. 1A-C). It distinguishes 17 different neuron types (listed in each row and column of Fig. 1B). These neuron types are further split into 111 different variations based on response profiles of individual neurons from the Allen Brain Atlas [Allen Institute, 2018], to which generalized leaky integrate-and-fire (GLIF_3_) neuron models with 3 internal variables had been fitted. These neuron models have in addition to the membrane potential two further internal variables that model after-spike currents (Fig. 1D). The resulting model for a patch of V1 receives visual input from an LGN model that consists of 2,589 filters [Billeh et al., 2020] that had been fitted to experimental data. This LGN model produces input currents to neurons of the V1 model in a retinotopic and lamina-specific manner [Billeh et al., 2020]. We employed a data-driven noise model based on experimental data from area V1 of the awake mouse [Stringer et al., 2019]. This noise model was not present in [Billeh et al., 2020] and had subsequently been introduced in [Chen et al., 2022]. It models both quickly changing forms of noise and slower forms of noise that contribute to experimentally found trial-to-trial variability (Methods).

**Figure 1:**
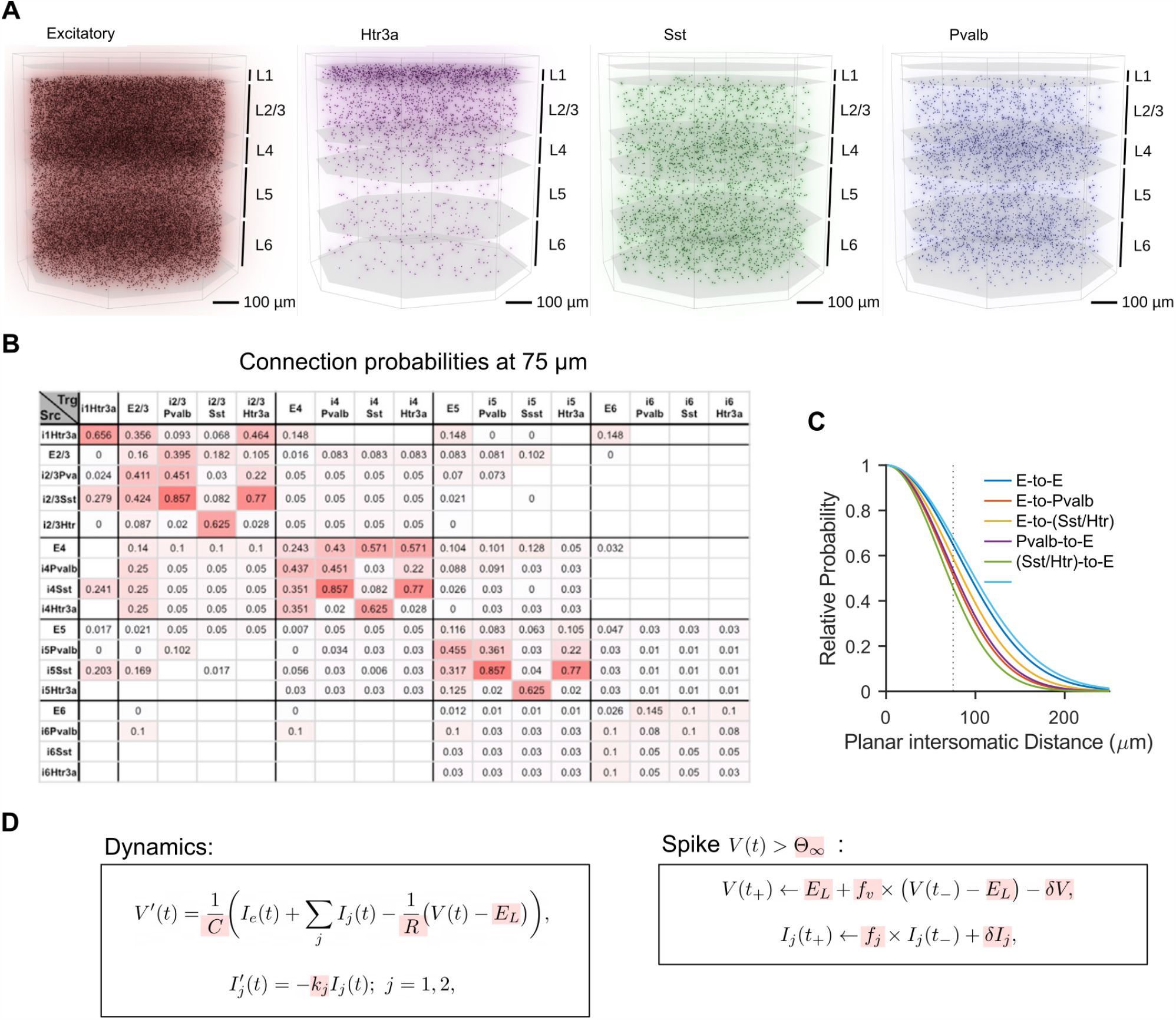
Structure and components of the data-based cortical microcircuit model of [Billeh et al., 2020]. **(A)** Side view of the 3D architecture of the model, which consists of with 51,978 neurons from 1 excitatory and 3 inhibitory neuron classes (Htr3a, Sst, Pvalb) in a column with 800 μm diameter. These neurons are distributed over 5 laminar sheets (L1 to L6, where L2 and L3 are lumped together). **(B)** The data-based base connection probabilities of [Billeh et al., 2020] depend on the cell class to which the presynaptic (row labels) and postsynaptic neuron (column labels) belongs. White grid cells denote unknown values. **(C)** The base connection probability from **(C)** is multiplied according to [Billeh et al., 2020]) for any given pair of neurons by an exponentially decaying factor that depends on the lateral distance between them. **(D)** Main equations defining the GLIF3 neuron models of [Billeh et al., 2020] with 3 internal variables. Assignments to their parameters (highlighted in red) define the 111 neuron models of the networks, based on experimental data from the Allen Brain Atlas [Allen Institute, 2018].

The resulting V1 model was trained to solve two computational tasks that have frequently been used in mouse experiments (Methods). One of them is the visual change-detection task, where a sequence of natural images is presented, interleaved with periods without visual input. The subject has to report whenever the most recently presented image differed from the one before (Fig. 2A-C). We randomly selected a pool of 140 natural images from the ImageNet dataset [Deng et al., 2009] that we used as network inputs. We used 40 of them for training, similar to the biological experiments of [Garrett et al., 2020b, Siegle et al., 2021], and 100 of them for testing (Fig. 2D). We compare details of the resulting network computation in the model with *in vivo* recordings from the visual cortex of the mouse for the same computational task. The other computational tasks that we considered is the evidence accumulation task (Fig. S1) that has been used in a fair number of mouse experiments [Driscoll et al., 2017, Morcos and Harvey, 2016b, Morcos and Harvey, 2016a, Koay et al., 2022], in which a mouse virtually navigates a linear track with visual cues, ultimately making a decision at a T-junction based on accumulated cues (Methods).

**Figure 2:**
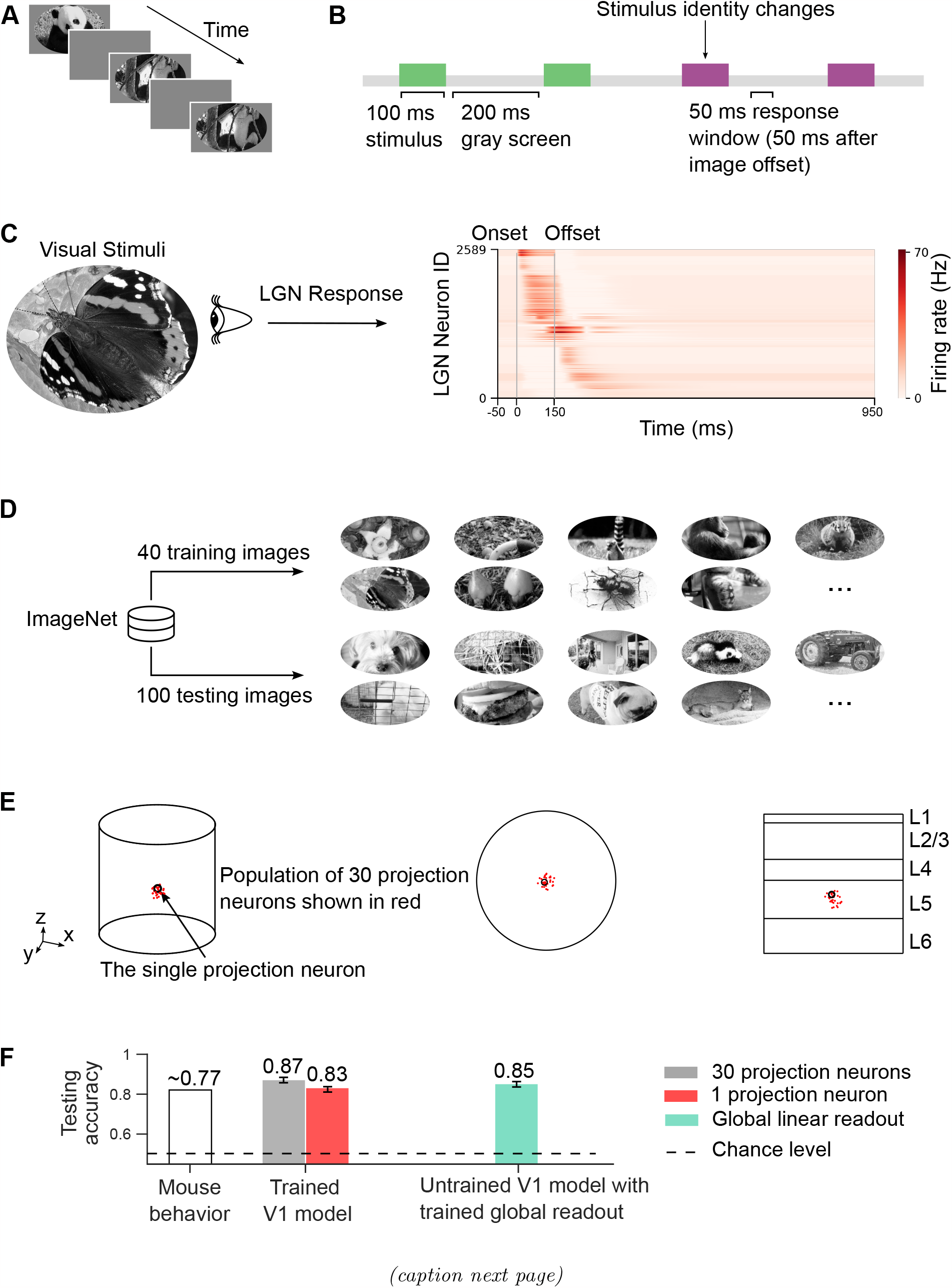
Visual-change-detection task. **(A)** A sequence of natural images is presented, interleaved with gray screen periods. **(B)** The visual-change-detection task requires to give during a 50 ms response window that begins 50 ms after image offset an output signal whenever the current image is different from the preceding one. **(C)** We used the LGN model from [Billeh et al., 2020] to transform visual stimuli into the outputs of 2,589 filters that model firing rates of LGN neurons and are connected to neurons in the V1 model according to data-based rules. These outputs provide input currents to V1 neurons using data-based rules. **(D)** We used 40 natural images for training and a separate set of 100 natural images for testing. Images were drawn from ImageNet [Deng et al., 2009] and presented in grayscale because of the insensitivity of LGN model to color. To ensure the robustness of our findings, we trained 10 V1 models using 10 different sets of training data (each with its own 40-image set). We found that all results presented below were consistent across all models, demonstrating that our conclusions are not reliant on specific training datasets or coincidences. **(E)** The network was trained to provide the network decision through projection neurons within the network, either a single (black circle) or 30 (red dots) randomly selected pyramidal cells in layer 5 (within a sphere of 55 μm) depending on the experiment. Their task was to report an image change through an increased sum of firing rates during the response window. **(F)** Comparable to the mouse behavior (left), task performance on the novel image set of the trained V1 model for reporting network decisions through increased firing activity of 1 or 30 projection neurons is shown in the middle. The number of projection neurons has little impact on the network performance. The green bar on the right shows a control result for the case when one uses instead of projection neurons from within the network a global linear readout from all neurons in the network. One sees that it is not even necessary to train the V1 model for the task: Training of the weights of the artificial global linear readout suffices, indicating that the computation within the V1 model can be effectively masked by such a global linear readout. The error bars, which are small (*<* 0.01), represent the standard error of the mean (SEM) across 10 models with different training datasets.

Pyramidal neurons on layer 5 of cortical microcircuits extract computational results of the microcircuit and transmit them to other brain areas, thereby triggering behavioral responses [Harris and Shepherd, 2015, Marshel et al., 2019]. Therefore we selected a set of pyramidal cells on layer 5 as projection neurons of the V1 model (Fig. 2E). These projection neurons had the task of reporting the network decision through firing during a specific response window. The size and spatial distribution of these projection neurons had little impact on the results (Fig. 2F). For simplicity, we used a single projection neuron, unless stated otherwise.

A more common output convention in modeling brain computations is to use an external readout neuron that receives synaptic input from all network neurons. We found that this convention is not suitable for probing the computational capability of a network model. First, such global readout neurons that receive synaptic inputs from all neurons in a large patch of the neocortex have not been found in the brain. Secondly, a global linear readout neuron that receives synaptic inputs from a large set of neurons in a recurrent neural network through trained synaptic weights is a too-powerful device that masks the computational contribution of the recurrent neural network. Instead, the recurrent neural network plays in this case just the role of a liquid or reservoir [Maass et al., 2002, Maass and Markram, 2004]. Concretely, if one takes the V1 and LGN model as defined in [Billeh et al., 2020], without changing any of its synaptic weights or other parameters, and just trains a linear readout from all of V1 neurons for the visual-change-detection task, one gets already a very high average accuracy of 0.85 (see the rightmost bar in Fig. 2F).

We trained all synaptic weights from the LGN to V1 and within the V1 model using stochastic gradient descent for a suitable loss function that assumed a low value only when the projection neuron(s) fired within a short response window 50 ms after the presentation of an image in case that this image differed from the preceding one (Methods). We included regularization terms similarly as in [Chen et al., 2022] in the loss function in order to keep the firing activity of the network in a biologically realistic sparse firing regime. Synaptic connectivity was not changed through this training process. We also did not allow synaptic weights to change their sign, thereby obeying Dale’s law. In particular, the average firing rate after training was around 3.9 Hz. Hence the model computed in an energy-efficient sparse firing regime. The new values of synaptic weights after training remained in a biologically reasonable range, see Fig. S2 and S3.

The trained V1 model achieved high performance for both tasks, which lies in the same range as the performance achieved by mice [Garrett et al., 2020b, Morcos and Harvey, 2016a]. In addition, the model was also able to generalize well for the visual-change-detection task, achieving almost the same performance for images that were not used during training (Fig. 2F). Hence the model had acquired a network algorithm that generalized, i.e., was not constrained to a particular set of previously seen images.

### 2.2 Information content of soft rank order coding in the V1 model

Simultaneous recordings from large numbers of neurons in the awake brain show that neural networks of the neocortex exhibit a peculiar type of network dynamics that is rarely seen in artificial neural networks: Most neurons fire only during a rather short time window during a trial, see e.g. [Driscoll et al., 2017] and Fig. 2 of [Koay et al., 2022]. Furthermore, different neurons have different preferred firing times, and the relative order in which they fire depends on the task condition and the sensory input. Hence, similarly as in synfire chain models [Abeles et al., 2004], the network activity has a prominent sequential character, but in contrast to synfire chain models, only a very small fraction of neurons is active during each segment of the sequence. In addition, the particular order of this sequential firing activity encodes computationally relevant information [Driscoll et al., 2017, Koay et al., 2022]. Recent theoretical work suggests the soft rank order coding of the neocortex is a close to optimal coding scheme from the perspective of energy consumption of 3D network computations [Boahen, 2022].

Soft rank order coding is reminiscent of previously considered coding schemes for spiking neurons, where the rank order of single spikes from different neurons on the time scale of millisecond was postulated to contain salient information [Thorpe, 1990, Maass, 1994]. But that scheme was hard to reconcile with experimental data, where neurons tend to fire a single spike in one trial, and several spikes or no spike in other trials. The experimentally found soft rank order coding takes place on the time scale of several hundreds of ms, i.e., on the time scale of behavior, and allows neurons to fire several spikes during the relevant time period. Therefore we refer to this coding method as soft rank order coding, rather than rank order coding.

Since largely sequential network activity is hard to reproduce in neural network models, we wondered whether the data-based V1 model would be able to do that. We considered trial-averaged neural activity as in [Koay et al., 2022] and followed the same routine to order the neurons according to the time of the peak activity for the visual-change-detection task. Furthermore, as in their data analysis, we normalized the firing activity, averaged over 200 trials, of each neuron over the 300 ms of the computation on each image, consisting of a segment of the continuous inputand processing stream including 100-ms image presentation, 150-ms delay, and 50-ms response window, marked at the top of Fig. 3 A-C. We found that the neural activity in the V1 model was indeed very similar to that in the experimental data. Fig. 3 A-C show that most neurons of the V1 model participated in the network computation, but focused their firing activity to a very short segment of the 300-ms time window. In addition, as in the experimental data, the relative order of these short segments of high activity depended on whether the currently processed image was the same as the preceding one or not, see Fig. 3. Panel A, where we have ordered the neurons according to their preferred firing time for the change condition, shows that the neurons have for the change condition a clear order of preferred firing times. A clear order is also shown in panel B for nochange condition. But the order is different from that for the change condition, as panel C shows: The resulting sequence is blurred and lacks the characteristic thin-line pattern observed in Fig. 3A and B. In other words, the network employed soft rank order coding through the relative timing of the peak-firing activity of neurons for distinguishing between these two experimental conditions.

**Figure 3:**
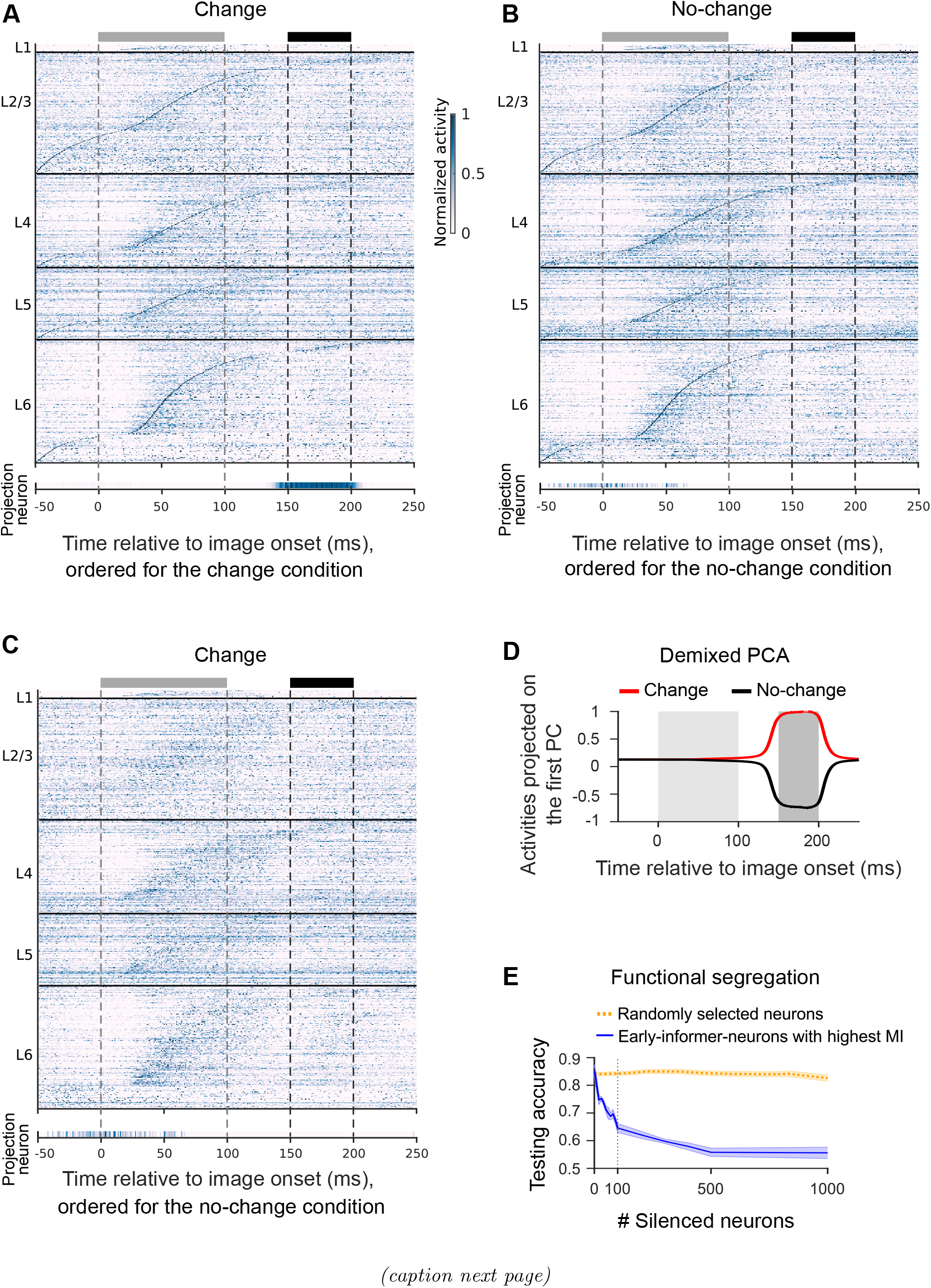
Temporal organization of computations in the V1 model. **(A)** As in the experimental data, neural activity is sparse and exhibits a clear sequential organization with high temporal resolution. Normalized average responses over 200 trials with the change condition but different images are shown, with neurons ordered according to the time of their peak activity under the change condition. The gray and black bars at the top denote the image presentation and response windows, respectively. **(B)** Same as in **(A)**, but for the no-change condition, with neurons ordered according to the time of their peak activity for the no-change condition. **(C)** The same data as in **(A)** for the change condition, but with neurons ordered as in **(B)**. The resulting blurred sequence indicates that the order of peak activity of neurons is quite different for the change and no-change conditions. **(D)** In order to visualize the formation of the network decision, we carried out demixed principal component analysis for trial-averaged network activity (Methods). Its projection onto the first principal component is shown. The network decision starts to emerge during the time window from 50 to 100 ms after image onset. The light and dark gray rectangles denote the window of image presentation and response, respectively. **(E)** Causal impact of specific neurons on the network decision: Task performance quickly decreases when early-informer-neurons are silenced (in the descending order of their MI with the network decision), see the blue curve. On the other hand, task performance is robust to silencing the same number of randomly selected neurons (dotted yellow curve). Both curves show average values for 10 V1 models where different sets of training data were used. The shaded area represents the SEM across 10 models.

We employ a statistical measure, Kendall’s *τ* coefficient between rank orders under change and no-change conditions, for quantifying this information (Methods). *τ* = 1 indicates that two orders are in complete agreement, signifying a perfect positive association. *τ* = *−*1 denotes a perfect negative association. *τ* = 0 means two orders are independent of each other. *τ* between orders under change and no-change conditions is near zero, *τ* = 0.004 *±* 0.001 (mean *±*standard deviation, calculated across 10 models each trained with distinct training data sets), indicating a significant disparity. Furthermore, this soft rank order coding was simultaneously expressed in all layers of the laminar cortical microcircuit model. Note that this coding type, through the order to peak activity of different neurons, takes place in spite of substantial noise both in the brain and in our V1 model, see [Chen et al., 2022] for details, that substantially affects the timing of individual spikes in a single trial. The V1 model predicts that cortical microcircuits employ an even more refined type of soft rank order coding: The order of peak activity also varies with the identity of the current image (Fig. S4) and the preceding image (Fig. S5). Kendall’s *τ* coefficients presented in Fig. 4B validate the distinction in sequential patterns across varying stimulus conditions (*τ* is calculated based on 10 novel images). We are not aware of previous experimental data from the mouse that have tested this prediction from the model: That the soft rank order of firing encodes also information about sensory stimuli. Therefore we have re-analyzed *in vivo* data, as reported in the next subsection.

**Figure 4:**
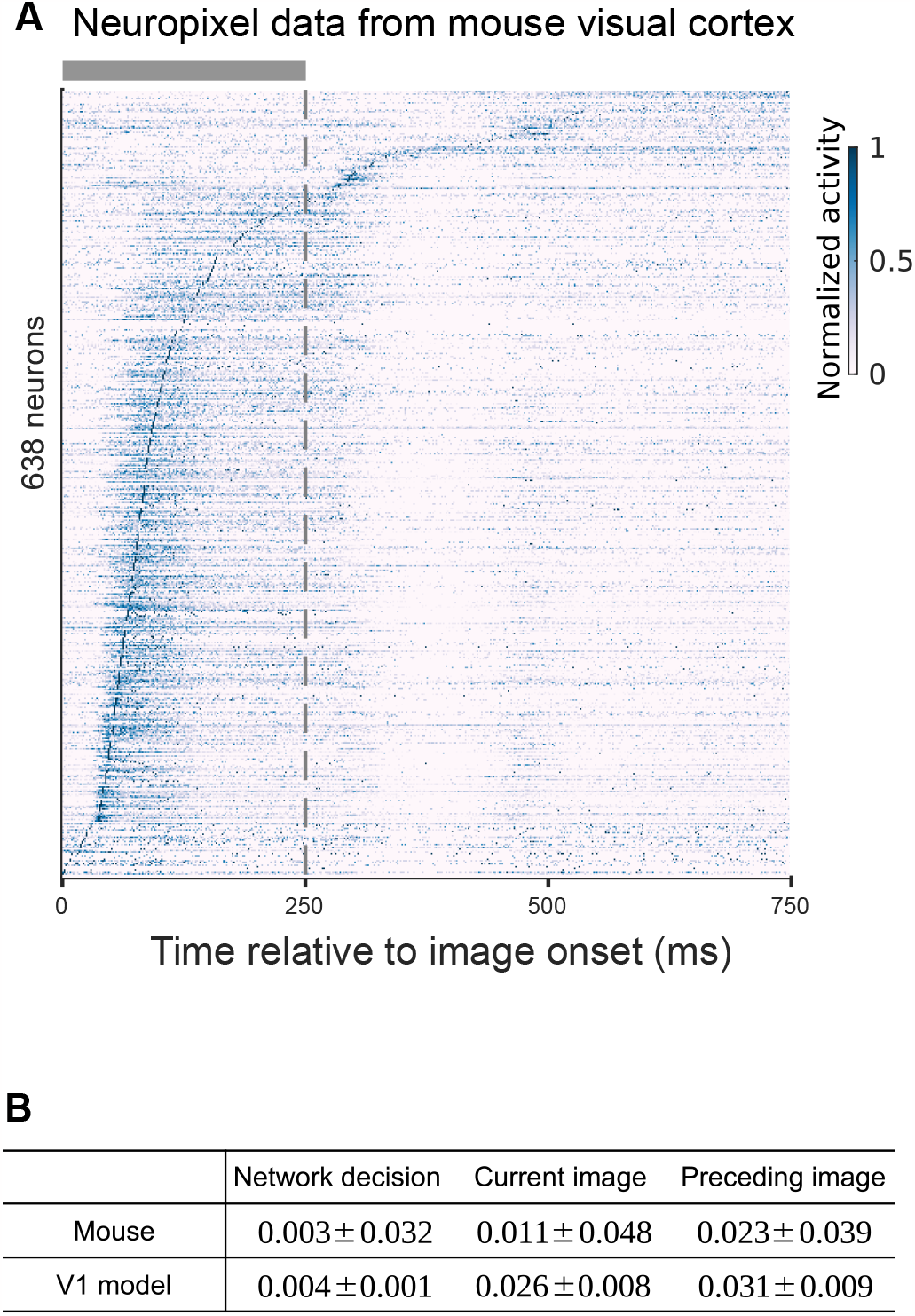
Test of the prediction of the V1 model regarding information content of soft rank order coding in V1 through re-analysis of *in vivo* neuropixel data. **(A)** Normalized average responses to novel images over 200 change trials from all recorded neurons in the visual cortex of mouse 548720 (ID in [Allen Institute MindScope Program, 2022]), which had been trained on the visual-change-detection task. Neurons are ordered by their peak activity time under the change condition. The gray bar indicates the 250-ms image presentation period, followed immediately by the response window. **(B)** Measurement of the content of information in the soft rank order code, using Kendall’s *τ*, about the network. Values are averaged across 20 mice (mean*±*standard deviation) that had all been trained on the visual-change-detection task. For comparison, we show corresponding measurements for the V1 model, averaged across 10 models trained with different training sets (mean*±*standard deviation). All values are close to 0, suggesting substantial information content in soft rank order coding. The values for the current and preceding image are closer to 0 for the *in vivo* data from mouse V1, suggesting that the model slightly underestimates this information content.

We have also analyzed in the V1 model the occurrence and information coding of soft rank order coding for the evidence accumulation task. We found that the network computations were organized into similar sequential patterns (Fig. S6A-C, Supplementary Note 1). The rank orders of neuronal peak activities for turning left and turning right were distinguishable, as evidenced by Kendall’s *τ* coefficient, *−*0.002 *±* 0.006, being close to 0.

### 2.3 Information content of soft rank order coding in mouse V1: Re-analysis of *in vivo* neuropixel data

We conducted an analysis of the sequential patterns in the mouse visual cortex during the visual-change-detection task (Methods), as recorded through neuropixels [Allen Institute MindScope Program, 2022]. While we examined responses to both familiar and novel images, we report only on the latter, mirroring our approach with the V1 model. Notably, the neuropixel data did not reveal significant differences between the responses to familiar versus novel images. In Fig. 4A, the neural activity prominently demonstrates sequential characteristics, with only a small fraction of neurons being active during each phase of the trial. This sequential firing pattern was evident across all subjects (*n* = 20 mice), and corroborates the sequential pattern identified in the V1 model (Fig. 3A). Moreover, we found that the sequential pattern in the mouse visual cortex not only encodes information about the network decision, but also about the current and preceding image. This finding is quantitatively validated by measurements of Kendall’s *τ* coefficients (Fig. 4B). Hence soft rank order coding in area V1 of the mouse does in fact contain information about visual stimuli, as predicted by the V1 model.

The dependence of soft rank order on the identity of a visual stimulus has recently already been reported for area V1 of the monkey, see Suppl. Fig. 14 of [Yiling et al., 2023]. The visual stimuli used there were somewhat different: 3 familiar natural images that were ramped up over a period of 2000 ms. Similar results are reported there for area V4 of the monkey, also for short or no ramping up of visual stimuli. In contrast, natural images were flashed on in the experiments of [Siegle et al., 2021] for the visual-change detection task and in our V1 model. Another difference is that we measured Kendall’s *τ* for a set of 10 novel natural images. We are not aware of prior experimental results that demonstrate that soft rank order codes contain not only information about the current image, but simultaneously also about the preceding image (while another image is currently shown), as we found in mouse V1.

### 2.4 Functional segregation and integration in cortical microcircuits

The experimental data of [Houweling and Brecht, 2008, Doron et al., 2014, Marshel et al., 2019, Dalgleish et al., 2020, Doron et al., 2020] have elucidated another characteristic trait of cortical computations: The network computation is surprisingly sensitive to the activity of a tiny fraction of neurons of these very large networks, since stimulation of a few selected neurons could change the result of the network computation, i.e., the behavioral response. This trait can be seen as an instantiating of the principle of combined functional segregation and integration, which has been postulated to characterize a fundamental property of brain networks [Tononi et al., 1998]. Functional segregation refers here to functional specialization of individual neurons, and functional integration to the capability of the network to properly integrate the responses of neurons with diverse specializations. The high sensitivity of the neocortex, or in our model of the projection neurons, to the activity of particular neurons is surprising insofar as the network has to cope with a substantial amount of noise, see the analysis and resulting noise model of [Chen et al., 2022] that we also used in this study.

We silenced individual neurons to test the capability of the projection neuron. While this manipulation is difficult *in vivo*, it is easy to carry out in the model. The common method for evaluating functional segregation *in vivo* is to activate selected neurons, but this has the disadvantage that one does not control how many other neurons are made to fire through that. Like in the *in vivo* data on the neocortex, we found that the result of the network computation is not equally sensitive to the activity of all of the neurons, but that there are particular neurons that are pivotal for the network decision. For the visualchange-detection task. We focused on silencing early-informer-neurons, i.e., neurons that produced the first information on whether the current image was the same as the preceding one, well before the response window, while the current image was still processed. Demixed principal component analysis (Methods) in Fig. 3D shows that the first information about this arises during the time window from 50 to 100 ms after image onset, when the first information about the image reaches the V1 part of the model. Hence we defined early-informer-neurons as neurons whose spike output contained already during this time window substantial mutual information (MI, Methods) with the upcoming network decision. Fig. 3E shows that while the network is not very sensitive to silencing of randomly selected neurons, silencing of just 100 early-informer-neurons has a drastic impact on the network performance. Also, the number of projection neurons does not affect the conclusion of the lesion experiment (Fig. S7). Supplementary Note 1 and Fig. S6D demonstrate similar results in evidence accumulation task.

Note that the special role that early-informer neurons play for the network computations can be seen as a clever combination of functional segregation (specialization) of particular sets of neurons with some form of integration, since the results of the early-informer-neurons reach the chosen projection neurons on layer 5 that report the network decision. This concept had been introduced in order to characterize on an abstract mathematical level computational traits of brain computations [Tononi et al., 1998]. The concept of functional segregation has subsequently been used quite frequently in the analysis of experimental data on whole-brain computations. For example, it was applied in [Long et al., 2016] in the analysis of the impact of cooling particular areas of the human brain on speech performance. Similarly, we use this term here in the analysis of the impact of particular sets of neurons on the computational performance of the network. We introduced the following quantitative index for evaluating the functional segregation of early-informer neurons, i.e., their impact on the network decision: (*A*_random_ *− A*_MI_)*/A*_random_, where *A*_random_ denotes the task performance (testing accuracy) obtained after silencing 100 randomly selected neurons, while *A*_MI_ refers to the task performance achieved after silencing 100 neurons with the highest mutual information (MI) in the time window [50, 100] ms. The mean functional segregation index of V1 model is 0.21 for both tasks.

### 2.5 Previously considered models for cortical computation cannot reproduce these characteristic traits

Many models have been introduced to simulate cortical computation. Notably, the randomly connected recurrent networks of artificial neurons (RANN), the recurrent convolution neural network (RCNN), and the generic recurrent spiking neural network (RSNN) stand out as popular choices among them. Yet, as we will illustrate through detailed quantitative analyses, these previously proposed models fail to reproduce the characteristic traits under discussion, although they achieve high accuracy on both tasks. We ensured consistency across the models by maintaining an identical count of neurons and synapses with the V1 model (for the RCNN, a comparable neuron count was used). All models underwent training using the same tasks, inputs, and optimization techniques, as detailed in the Methods section.

RANNs have commonly been used as models for cortical computations [Sussillo and Barak, 2013, Sussillo et al., 2015, Yang et al., 2019, Yang and Wang, 2020, Pollock and Jazayeri, 2020]. We used a standard neuron model from RANN models for neural networks of the neocortex [Sussillo et al., 2015, Pollock and Jazayeri, 2020]: A non-spiking neuron with tanh as activation function and a membrane time constant of 50 ms. For extracting the network decision we used the same global linear readout from all neurons in the RANN as in these paradigms. This RANN model was after training able to perform the visualchange-detection task at a higher performance level than the data-based V1 model and the subjects in the neurobiological experiments [Garrett et al., 2020b]. However, Fig. S8 A and B show that neurons of the RANN do not have a similarly short time period of high activity during the visual-change-detection task. Also, the temporal order of neural activity in RANN does not depend on the trial type, see Fig. S8C. This is confirmed by a notably larger amplitude of Kendall’s *τ*, which is three orders of magnitude greater than that observed in the V1 model and mouse visual cortex (Table 1). Since the activity level of neurons in the RANN depends on the weight of the regularization term in the loss function for stochastic gradient descent, we repeated the training of the RANN with different weights of this regularization term that controls the average activity of neurons in the network (Fig. S9). Also in these controls, the neurons of the RANN do not constrain their activity to a short time window like in the experimental data and the V1 model. The same holds for a further control where we changed the threshold of the regularization term in the loss function (Methods). These results suggest that, under typical configurations, the RANN model can not naturally reproduce the sparse neural activity with trial-type-dependent temporal sequences of neural peak activity observed in the neocortex.

**Table 1:** Quantitative analyses for the visual-change-detection task: A comparison of task performance, soft rank order coding of the network decision (using Kendall’s *τ* coefficient), and the functional segregation index across mouse experiments and different models. For mice, *τ* was derived from neuropixel data of 20 mice. For models, *τ* and index of models were calculated across 10 models trained with different training sets, presented as mean*±*standard deviation.

| | Task performance<br>on novel images | Kendall’s $\tau$ | Functional<br>segregation index |
| --- | --- | --- | --- |
| Mouse | 0.77<sup>a</sup> | $0.003 \pm 0.032$ | Experimental evidence<sup>b</sup> |
| V1 model | 0.83 | $0.004 \pm 0.001$ | $0.21 \pm 0.02$ |
| V1 model without LGN<br>(control model 1) | 0.83 | $-0.009 \pm 0.004$ | $0.22 \pm 0.02$ |
| V1 model without neuron<br>diversity (control model 2) | 0.81 | $0.042 \pm 0.042$ | $0.07 \pm 0.02$ |
| V1 model without laminar<br>structure (control model 3) | 0.80 | $-0.023 \pm 0.082$ | $0.10 \pm 0.02$ |
| RSNN<br>(control model 4) | 0.79 | $0.243 \pm 0.012$ | $0.03 \pm 0.03$ |
| RSNN with E/I distinction<sup>c</sup><br>(control model 5) | 0.50 | N/A | N/A |
| RANN | 0.97 | $0.273 \pm 0.038$ | $0.02 \pm 0.02$ |
| RCNN | 0.97 | $0.273 \pm 0.082$ | $0.02 \pm 0.02$ |

(a) Estimated from Fig. 1I of [Garrett et al., 2020a] when novel images were presented to mice.
(b) Conclusion from [Houweling and Brecht, 2008, Doron et al., 2014, Marshel et al., 2019, Dalglish et al., 2020, Doron et al., 2020]: activation of very few neurons caused perceptual or behavioral changes.
(c) Control model 5 cannot solve the visual-change-detection task (chancel-level accuracy).

We also tested the functional segregation of the RANN and found that about 6,000 neurons need to be silenced in the RANN to reduce the accuracy of the network computation to 0.6 (Fig. S8D), a performance level which was reached in the V1 model according to Fig. 3D by silencing just 200 neurons. Note that we silenced here RANN neurons in descending order of the MI of their activity during the second half of an image presentation with the upcoming network decision, like in the V1 model. The RANN has more high-MI neurons than the V1 model (Fig. S10). A notably lower functional segregation index of 0.002 further underscores the significant divergence from the V1 model. These results suggest that the RANN is substantially less sensitive to the activity of small subsets of its neurons.

In Supplementary Note 2, we demonstrate that the RCNN and RSNN models also do not reproduce the two characteristic traits of cortical computation for the visual-change-detection task. The sequential patterns, as illustrated in Figs. S11 and S12, combined with the metrics in Table 1, reveal their deviation from both the experimental data and the V1 model. Also, Supplementary Note 1 addresses the inability of RANN (Fig. S13), RCNN, and RSNN to capture these traits also for the evidence accumulation task, supported by the quantitative data in Table 2.

**Table 2:** Quantitative analyses for the evidence accumulation task: A comparison of task performance, soft rank order coding of the network decision (using Kendall’s *τ* coefficient), and the functional segregation index across mouse experiments and different models.

| | Task performance | Kendall’s $\tau$ | Functional segregation index |
| --- | --- | --- | --- |
| Mouse | $\sim 0.85^a$ | N/A | Experimental evidence<sup>c</sup> |
| V1 model | 0.83 | $-0.002 \pm 0.006$ | $0.21 \pm 0.01$ |
| V1 model without LGN<br>(control model 1) | 0.83 | $0.003 \pm 0.004$ | $0.21 \pm 0.01$ |
| V1 model without neuron<br>diversity (control model 2) | 0.81 | $0.058 \pm 0.018$ | $0.08 \pm 0.02$ |
| V1 model without laminar<br>structure (control model 3) | 0.80 | $-0.023 \pm 0.056$ | $0.11 \pm 0.03$ |
| RSNN<br>(control model 4)<sup>c</sup> | 0.50 | N/A | N/A |
| RSNN with E/I distinction<br>(control model 5)<sup>c</sup> | 0.50 | N/A | N/A |
| RANN | 0.97 | $0.273 \pm 0.036$ | $0.02 \pm 0.02$ |
| RCNN | 0.97 | $0.263 \pm 0.094$ | $0.02 \pm 0.02$ |

(a) Estimated from Fig. 1C of [Morcos and Harvey, 2016a] when the number of cues was 6.
(b) Conclusion from [Houweling and Brecht, 2008, Doron et al., 2014, Marshel et al., 2019, Dalglish et al., 2020, Doron et al., 2020]: activation of very few neurons caused perceptual or behavioral changes.
(c) Control model 4 and 5 cannot solve the evidence accumulation task (chance-level accuracy).

The RSNN with excitatory and inhibitory neurons (control model 5 as defined later) plays a special role since it could not be trained by BPTT to exceed the chance level in performing the visual-change detection task (or the evidence accumulation task). However, this model exhibits according to [Yang and Wang, 2020] a stimulus-dependent rank order coding, although only over 30 ms for flashed-on stimuli. Hence its neural coding and computational properties are also very different from those of the V1 model.

### 2.6 Identification of causal relations between features of the V1 model and the emergence of characteristic traits of cortical computation

To understand the critical attributes necessary for reproducing the unique aspects of cortical computation, we next embark on quantitatively comparative analyses of the V1 model against five control models (Methods, Table S1), each differing in specific biological features. In control model 1, we removed the LGN model of [Billeh et al., 2020], and injected pixel values directly into the V1 model, targeting the same neurons as the LGN model. In control model 2, we removed the diversity of the 111 data-based neuron types in the V1 model, replacing them with one generic model for excitatory neurons and one for inhibitory neurons. In control model 3, we removed instead the laminar spatial structure with distance-dependent connection probabilities of the V1 model, replacing them with an equal number of randomly chosen connections (without dependence on spatial proximity). Control model 4 is RSNN (same as in Sec. 2.5), with the same number of neurons and connections. Control model 5 is a variation of it, where the neurons are split like in the V1 model into excitatory and inhibitory neurons, and Dale’s law is preserved during training.

As demonstrated in Table 1 and 2, we present the performance metrics, Kendall’s *τ*, and functional segregation index for the V1 model alongside the control models to quantify the changes of two notable characteristics. The influence of the LGN model is minimal (as seen in control model 1), aligning closely with the V1 model. In contrast, the impacts of neuron diversity and laminar structure are considerably more pronounced (evident in control models 2 and 3). Here, the absolute value of Kendall’s *τ* is approximately tenfold that of the V1 model, and the functional segregation index is roughly a third. The RSNN, designated as control model 4, while frequently used to emulate the cortex, lacks many biological features. Its behavior diverges significantly from that of the V1 model, with a three-order magnitude difference in Kendall’s *τ* and the lowest sensitivity index. Control model 5 fails to solve the visual-change-detection task. Taken together, our quantitative analyses indicate that neuron diversity and laminar structure play pivotal roles in shaping the two characteristic traits of cortical computations under study.

### 2.7 Computational progress of the V1 model becomes visible as nested bifurcations of the network dynamics

Analyses of results of simultaneous recordings from large numbers of neurons in the brain have shown that low-dimensional projections of the high-dimensional network activity provide interesting links between the network dynamics and the computations that it performs [Broome et al., 2006, Kato et al., 2015, Allen et al., 2017, Steinmetz et al., 2019]. We wondered whether similar analyses could elucidate computations of the V1 model. We embedded its activity vectors in the visual-change-detection task into 2 dimensions by UMAP (Methods). The processing of each particular image gives rise to a bundle of trajectories, with trial-to-trial variability resulting from the preceding images and noise within the network. Two nested bifurcations in these bundles of trajectories mark the computational progress of the dynamical system, see Fig. 5A. First, the trajectories of network states bifurcate during the first 50 ms of an image presentation from the mid-region of the plotted state space and move into a region that is characteristic for the identity of the currently presented image, see Fig. 5B. Afterward, between 50 and 100 ms after image onset, the second bifurcation occurs in dependence on whether the current image is the same or different from the previously presented image, see Fig. 5C. These modeling results provide concrete predictions for the way how these computations are carried out by cortical microcircuits of the brain, viewed as dynamical systems. Among various options on how these could compute, as discussed in [Rabinovich et al., 2006], nested bifurcations of bundles of trajectories emerge as the clearest visible fingerprint of these network computations. Supplementary Note 2 and Fig. S14 demonstrate the trajectories for evidence accumulation task.

**Figure 5:**
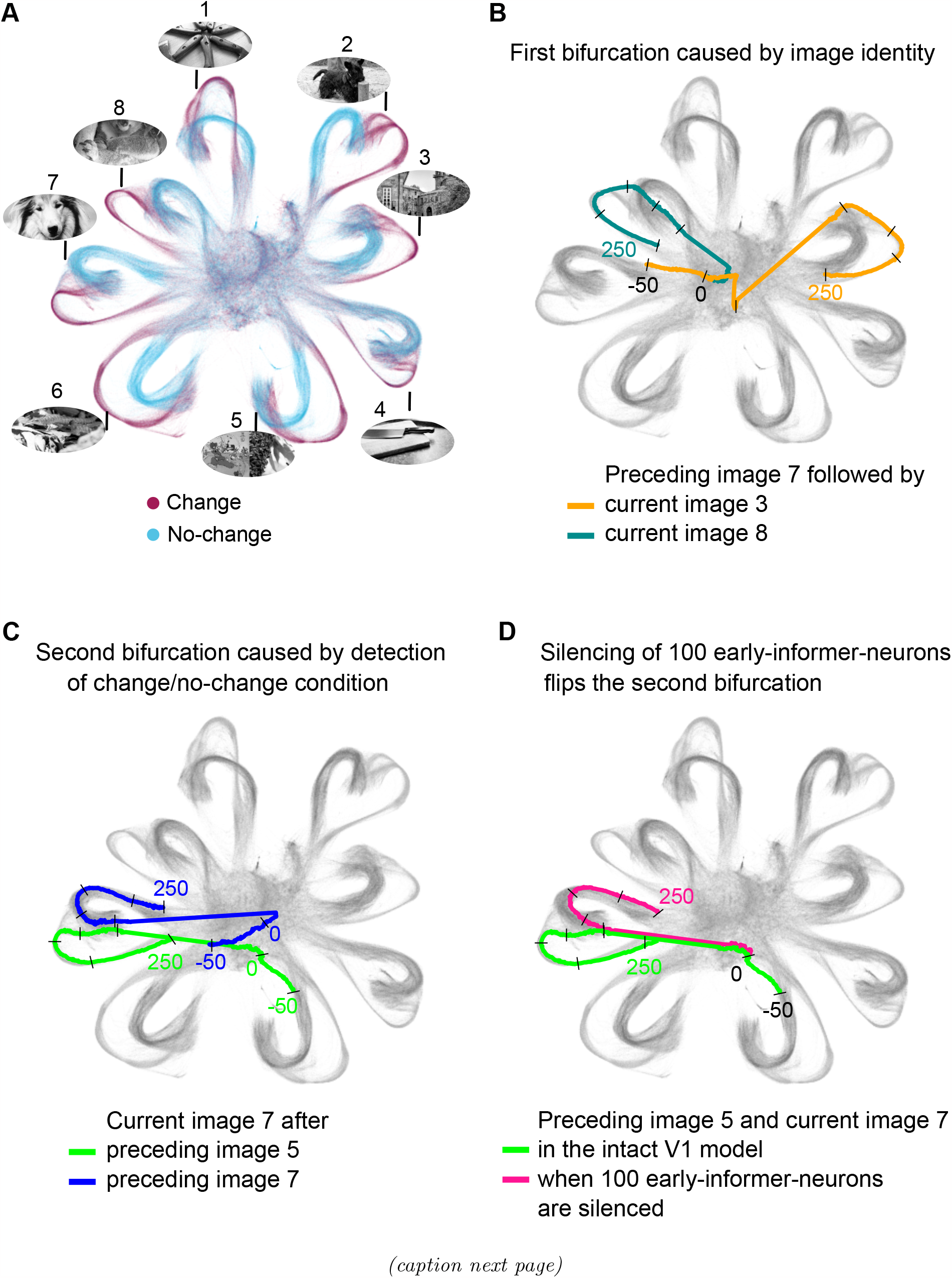
Nested bifurcations of trajectories of network states provide links between network dynamics and network computations. Each dot represents network activity at a specific millisecond within a 300-ms fragment of the computation of the V1 model on a sequence of natural images. Short black bars mark 50-ms long subsections of network trajectories. **(A)** Network trajectories of the V1 model during the presentation of 8 (out of the 100) test images that had not been presented during training. Two colors indicate whether the current image was identical (blue) or different (brown) from the preceding image. One sees that the network state moves for each image into a different region of the state space, no matter whether it was the same or different from the preceding image. **(B)** The first bifurcation occurs according to the identity of the current image, highlighted here for the case where the trajectory starts in both cases from the same state, which largely results from the identity of the preceding image. This first bifurcation occurs within the first 50 ms after image onset. **(C)** The second bifurcation does not depend on the identity of the current image, but on whether it is the same or different from the preceding image. The second bifurcation is more difficult to visualize since the trajectory arrives from two different regions that are characteristic of the identity of the preceding image. **(D)** A small set of early-informer-neurons is causal for the second bifurcation. Here image 7 is shown after image 5, both in the intact model and when 100 early-informer neurons are silenced. One clearly sees that these neurons are causal for the second bifurcation occurring within 50 to 100 ms after image onset, since silencing them lets the trajectory flip to the bundle for the no-change condition. Silencing of these neurons also flips a trajectory for the no-change condition to the bundle for the change condition, see Fig. S15. Note that the displayed altered trajectories represent trials in which the network made incorrect decisions.

### 2.8 Causal relations between the activity of individual neurons and the network decision

The result of our lesion experiments in Fig. 3E indicates that the firing activity of a specific small set of neurons was causal for the network decision. These early-informer-neurons were distinguished by the fact that their firing activity during 50 to 100 ms after image onset contained substantial MI with the upcoming network decision. We show in Fig. 5D that their firing activity in the visual-change-detection task was also pivotal for the network dynamics. Silencing the 100 early-informer-neurons with the highest mutual information (MI) with the upcoming network decision flips the trajectory of the network dynamics at the 2nd bifurcation to another bundle, see the magenta curve in Fig. 5D. This suggests that these bundles of trajectories had a certain attractor quality for time-varying network states. The model received for both trajectories shown in this panel exactly the same network input, hence differences were only caused by the silencing of the 100 early-informer-neurons (with some further variance possibly caused by ongoing noise in the network). This was a change trial, as the green curve in Fig. 5D indicates for the intact network (compared with the trajectory bundles in Fig. 5A). Silencing of these 100 neurons can also flip the trajectory of a no-change trial to the bundle of trajectories for change trials, see Fig. S15.

We then investigated what mechanism enabled these 100 early-informer-neurons to decide whether the currently presented image differed from the previously presented one. These neurons were primarily located in layers 2/3 and 4, see Fig. 6A. We selected 7 of them (Methods) from the 4 major neuron classes so that these had during the interval from 50 to 100 ms after image onset the largest MI with the subsequent network decision. Their firing activity is shown in Fig. 6B. One sees that they fired at different rates during the interval from 50 to 100 ms after image onset for the change and no-change conditions. These rate differences continued to be present during the subsequent delay and response window.

**Figure 6:**
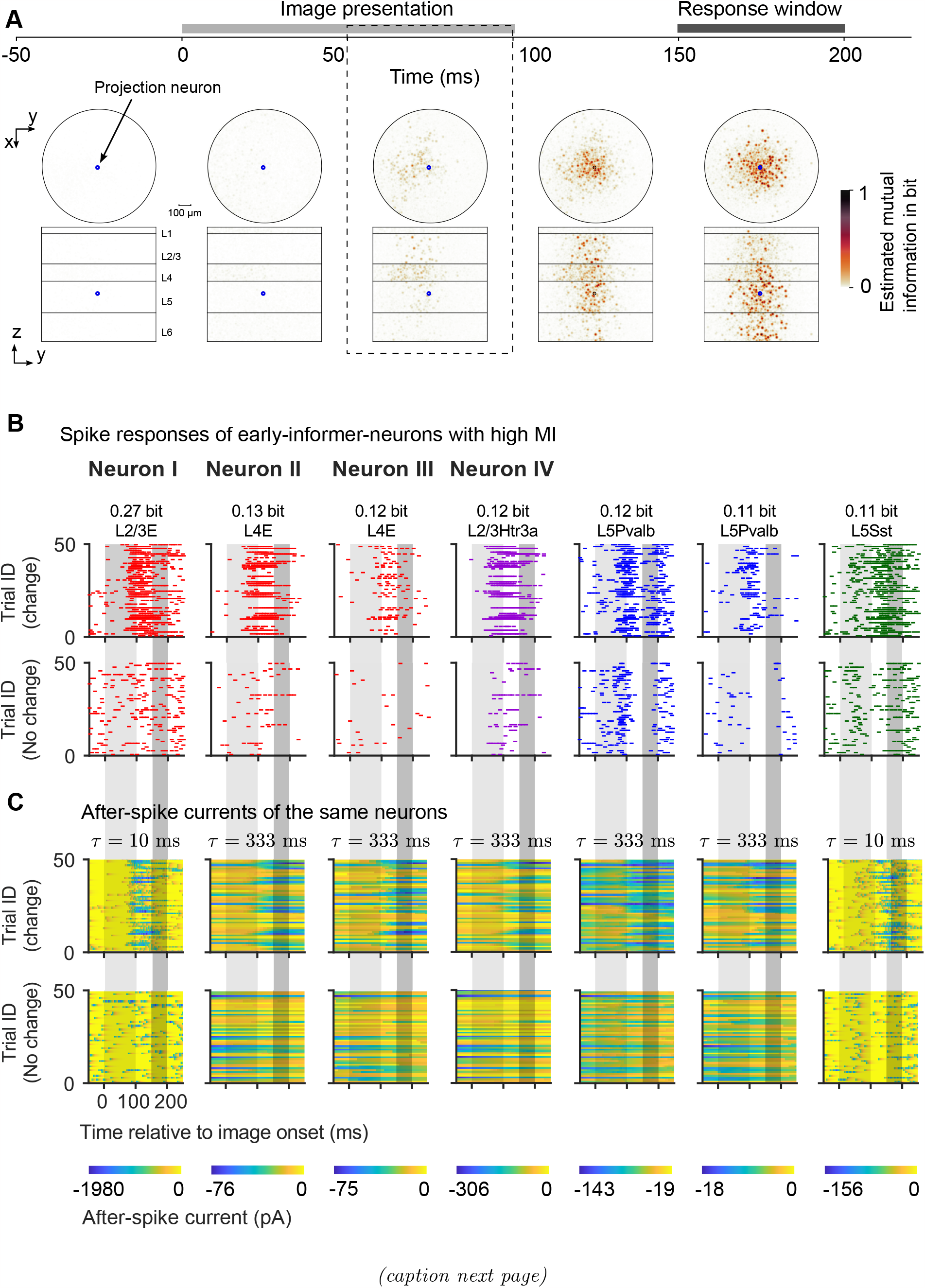
Fine-scale analysis of the emergence of a network decision in the V1 model. **(A)** The mutual information (MI) between activities in 50 ms windows of single neurons and the change/no-change decision of the network (as emerging during the subsequent response window) is estimated for each neuron. For neurons that overlap in the projection from 3D to 2D, the maximum value is visualized to avoid dark points arising from accumulation of small contributions from several neurons. The critical time period from 50 to 100 ms after image onset is marked by dashed lines. **(B)** Spike responses of 7 early-informer-neurons, selected to cover all four basic neuron types: excitatory (E), Htra3, Sst, and Pvalb neurons (color-coded as in Fig. 1A). L2/3, L4, and L5 represent layer 2/3, 4, and 5, respectively. The MI of the firing activity of each of these neurons during [50, 100] ms with the network decision is indicated in bits at the top of each panel. The periods of image presentation and subsequent response window are shaded in grey. Trials are separated according to the change/no-change condition. Condition-dependent differences in the firing responses of these neurons first appear at the start of the critical time window, 50 ms after image onset. **(C)** Time course of the internal variables of these neurons that had the longest time constant (after-spike currents), for the same trials as in **(B)** (time constant shown on top). As GLIF_3_ neurons have two after-spike currents that could have different time constants, we only show those with the largest time constant. Their values at image onset could potentially contain information about the identity of the preceding image, shown 200 ms before the current one. This information will be analyzed in the next figure.

In order to determine the mechanism by which these neurons acquired their early information about the relationship between the identity of the current and preceding image we analyzed the dynamics of their internal variables. Fig. 6C depicts the time course of their internal variables with the largest time constant. One sees that in many trials for 5 of these neurons, this internal variable had already a strongly negative value at the beginning of the presentation of the current image (indicated by the beginning of the light-gray zone). In order to understand the computational role of these internal variables, we analyzed whether their strongly negative values at image onset provided information about the identity of the preceding image. The result of this analysis is plotted for the first 4 neurons, whose locations are marked in Fig. 7A, B. Among the four neurons, three (represented in columns 2 to 4) had an internal variable with a large time constant. For these three neurons, the slow internal variable reached its lowest values at the onset of an image if a specific image (number 5) had been shown previously. Note, however, that these neurons could not specialize in reporting this particular preceding image, because the network was tested on new test images that had not been shown during training. Therefore, in order to serve as early-informer-neurons, the image features that produced the lowest values of their internal variables at the beginning of the next image had to tile the image space. The neuron whose analysis is plotted in the first column has only short time constants, and the value of its internal variable is less characteristic for a particular preceding image. Hence it is likely to collect and transmit information that it receives from other early-informer-neurons. In order to further elucidate the causal relation between the network decision and preceding activity (or non-activity) of early-informer-neurons with and without long time constants, we separately silenced from each of these two neuron classes those which had the largest MI with the network decision. The result in Fig. 7C shows that the network performance is significantly reduced when we silence 100 neurons with long time constants, and more than 500 neurons with short time constants need to be silenced to produce a similar reduction of network performance.

**Figure 7:**
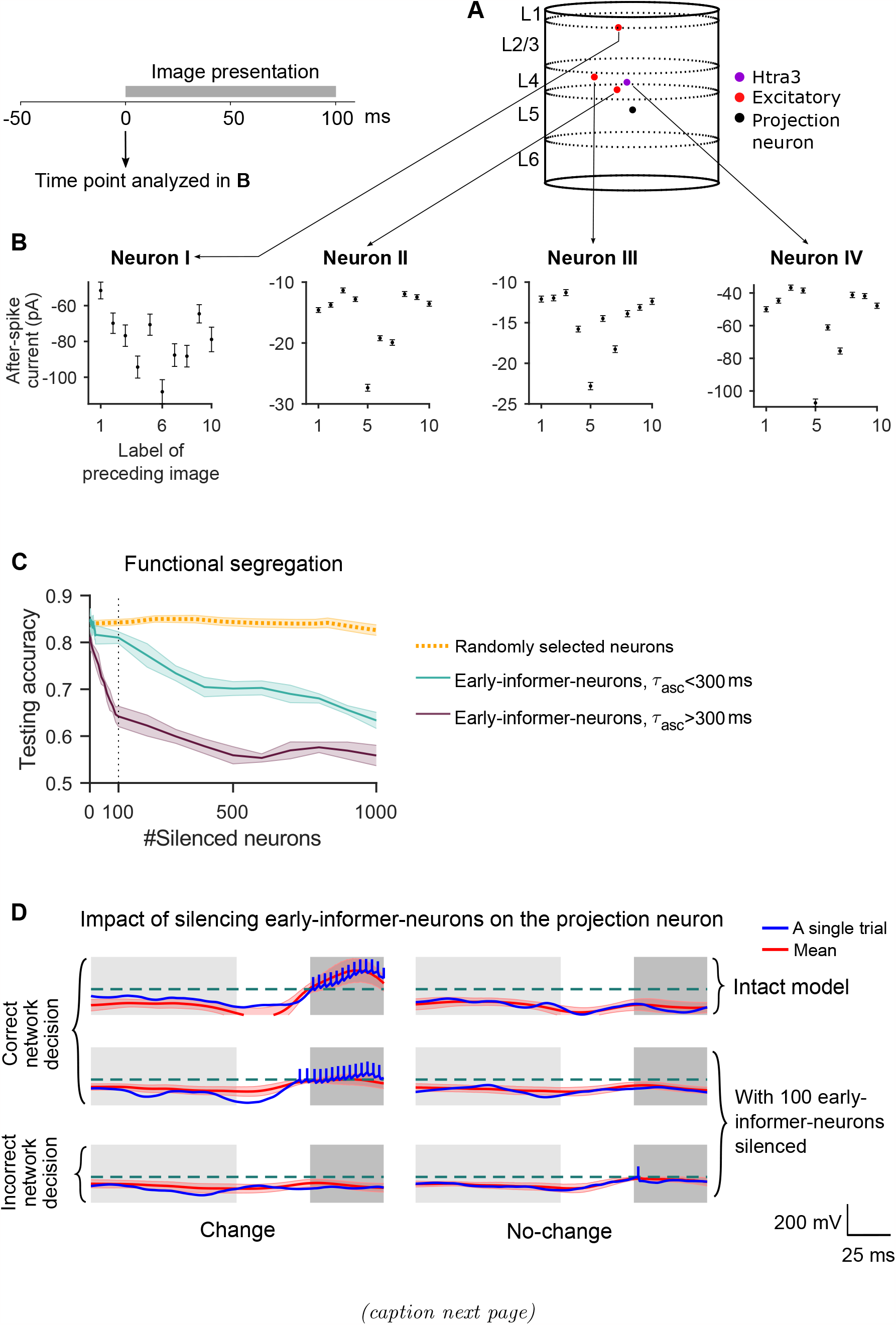
Inner workings of 4 sample neurons that are pivotal for the network decision. **(A)** Spatial locations and types of the 4 neurons labeled as Neuron I-IV in Fig. 6B and C are shown. Their firing rates during [50, 100] ms after image onset have the largest MI with the network decision that is made 100 ms later during the response window. **(B)** Analysis of the information contained in the after-spike currents of the four neurons. The after-spike current with the largest time constant is analyzed at the onset of the current image (see time point marked in the scheme above), depending on the identity of the PRECEDING test image, marked on the horizontal axis. Its mean value is shown with error bars denoting the SEM across 1000 trials. One sees that this value depends strongly on the identity of the preceding image, especially for neurons II - IV. **(C)** Silencing early-informer-neurons with after-spike currents that have large time constants (*τ*_asc_ *>* 300 ms) in the descending order of their MI reduces testing performance much faster than silencing those with shorter ones (*τ*_asc_ *<* 300 ms). The shaded areas represent the SEM across 10 models with different training datasets. **(D)** The membrane potential of the projection neuron is significantly changed under both the change and the no-change condition when 100 early-informer-neurons are silenced. We used the same stimuli but different realizations of the noise model and initial states. The dashed green lines represent the firing threshold. Blue curves represent the membrane potentials of single trials; the extended vertical bars denote the spikes in these trials. Red curves represent the average membrane potentials across 100 trials. The shaded red area represents the standard deviation across 100 trials.

The impact of early-informer-neurons on the firing or non-firing of the projection neuron was in general rather indirect, because they were in general not directly connected to the projection neuron (Fig. S16). This had to be expected, because the response window came 100 ms after the critical period of early-informer neurons (50 to 100 ms after image onset). Nevertheless, a direct impact of silencing the 100 early-informer-neurons on the membrane potential of the projection neuron can be detected. The result is shown in Fig. 7D. The 2nd and 3rd rows show that this silencing notably shifted the membrane potential of the projection neuron during the response window. This shift explain why their silencing led to errors in the network decision.

## 3 Discussion

Understanding the organization of brain computations, in particular computational processes in the neocortex, is one of the most exciting and pressing open problem in computational sciences. We have shown that a new method for advancing insight into the organization of cortical computations has now become feasible: We can combine insight from *in vivo* experiments with analyses of data-based cortical microcircuit models that are trained for the same computational tasks as the subjects of *in vivo* experiments. We have focused on two concrete visual processing tasks that have frequently been used for *in vivo* experiments: The visual change detection task and the evidence accumulation task. We demonstrated that a trained data-based microcircuit model can reproduce not only the task performance of *in vivo* subjects but also key traits of *in vivo* cortical computations, which are challenging to reproduce in generic recurrent networks, whether spiking or non-spiking. Furthermore, one can use the model to elucidate which features of cortical microcircuits are causally related to particular traits of coding and computational organization in cortical microcircuits. This is possible because one can modify the model, scramble for example its laminar architecture, or reduce the number of its different neuron types. These manipulations are currently not possible *in vivo*.

We have focused on the following two characteristic traits of *in vivo* computations that are hard to reproduce in generic spiking or non-spiking recurrent neural networks: Sparse sequential network dynamics that encoded information through soft rank order coding, and an exquisite combination of functional segregation and integration on the level of single neurons and microcircuits. The inability of artificial neural network models to readily reproduce these traits, despite performing the same tasks, highlights fundamental differences between the organization of cortical computations and that of artificial neural networks. Our control experiments have shown that both the laminar architecture of cortical microcircuits and their diversity of neuron types cause these differences in the organization of computations in cortical and artificial neural networks. We made this analysis rigorous by introducing quantitative measures for both of the two characteristic traits.

The data-based model allowed us also to take a look inside the neurons, and analyze which factors were essential for their functional segregation. We found that the values of internal variables with long time constants, which are abundantly present in the generalized leaky integrate-and-fire (GLIF_3_) neuron models of Billeh et al. (fitted to data of specific neurons from the Allen Brain Atlas [Allen Institute, 2018]) played an essential role for functional segregation in the visual change detection task: They assumed values at the beginning of the processing of the current image that contained substantial information about the identity of the preceding image (Fig. 6C and 7B). The specific causal impact of these neurons with long time constants on the network decision was verified through further lesion experiments in our model (Fig. 7D)

Since the same image-change detection task that we considered in the model was also used for *in vivo* experiments, we were able to test a hypothesis about the information content of soft rank order codes that have emerged both in the model and *in vivo*: Our analysis in the model showed that these soft rank order codes contain substantial information about the identity of the currently shown image, and also about the preceding images. We re-analyzed data from *in vivo* recordings from mouse V1 for the same task, and found that this also holds *in vivo*. This provides an example for the more direct interplay between models and *in vivo* experiments that becomes possible through the use of functional models that integrate most of our knowledge about the structure of cortical microcircuits. In particular, these models can produce predictions for the computational role of specific types and laminar locations of neurons that can be directly tested *in vivo*.

Another methodological advance is the measurement of the flow of information between specific types of neurons and their contribution to the network decision in the data-based model. This information-theoretic analysis can in general not be carried out *in vivo*, because it requires too many repetitions of the same trial under invariant conditions. But the predictions of this analysis can be tested through future *in vivo* experiments: The V1 model suggests that the network dynamics exhibits nested bifurcations, where each bifurcation is controlled by a rather small set of neurons that have collected particular pieces of information from the most recent sensory stimuli. The pivotal role of these small sets of neurons underlines the relevance of functional segregation of neurons for the network computations. Testing these model predictions *in vivo* will provide substantial progress in our quest to understand the organization of cortical computations.

Another next step in this quest for understanding the organization of brain computations will be to carry out similar analyses of the information flow, network dynamics, and neural coding in detailed models of cortical microcircuits from other species, and from other cortical areas, e.g. from the motor cortex. It is also feasible to tackle with the same method open questions about distributed computations in interconnected microcircuits from different cortical areas. Furthermore, to make the impact of top-down inputs more realistic, at least simplified models for the dendritic arborization of certain neuron classes should be incorporated. But this needs to be done in a way that still supports fast simulations of largescale models, so that training of these models for specific computational tasks remains computationally feasible. In addition, the computational role of projections to and synaptic inputs from subcortical areas, such as basal ganglia and the thalamus (see [Cruz et al., 2023] for a recent review), needs to be modeled and analyzed through integrated large-scale models. Altogether, this work is likely to complement, challenge, and enhance experimental work *in vivo* that aims at clarifying the organization of brain computations.

Our results, combined with corresponding *in vivo* data, suggest that coding and computing in cortical microcircuits are organized in a way that differs significantly from that in neural network models that are prevalent in current AI. Since cortical microcircuits are highly performant and noise-robust, and also operate in a substantially more energy-efficient sparse activity regime, our new research method and resulting insight into the relation between structure and function of cortical microcircuits is likely to impact future neural network models and computing technology in AI.

## 4 Methods

### 4.1 Neuron models

As in [Chen et al., 2022] we are focusing on the “core” part of the point-neuron version of the realistic V1 model introduced by [Billeh et al., 2020]. To make it gradient-friendly, we replaced the hard reset of membrane potential after a spike emerges with the reduction of membrane potential *z*_*j*_(*t*) (*v*_th_ *− E*_*L*_), where *z*_*j*_(*t*) = 1 when neuron *j* fires at time *t* and *z*_*j*_(*t*) = 0 otherwise. *v*_th_ is the firing threshold of membrane potential. *E*_*L*_ the resting membrane potential. This causes no significant change in the neural response [Chen et al., 2022]. We simulated each trial for 600 ms. The dynamics of the modified GLIF_3_ model was defined as

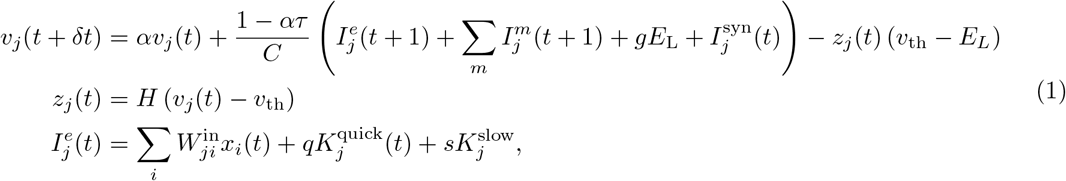

where *C* represents the neuron capacitance, *I*^*e*^ the external current, *I*^syn^ the synaptic current, *g* the membrane conductance, and *v*_th_ the spiking threshold. 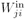 is the synaptic weight from LGN neuron *i* to V1 neuron *j*. The scales of the quick noise 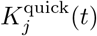 and the slow noise 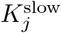 to neuron *j* are *q* = 2 and *s* = 2, respectively, unless otherwise stated. *K*_*j*_ was randomly drawn from the empirical noise distribution which will be elaborated on later. The decay factor *α* is given by *e*^*−δt/τ*^, where *τ* is the membrane time constant. *δt* denotes the discrete-time step size, which is set to 1 ms in our simulations. *H* denotes the Heaviside step function. To introduce a simple model of neuronal refractoriness, we further assumed that *z*_*j*_(*t*) is fixed to 0 after each spike of neuron *j* for a short refractory period depending on the neuron type. The after-spike current *I*^*m*^(*t*) was modeled as

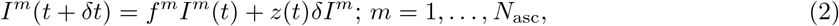

where the multiplicative constant *f*^*m*^ = exp ( − *k*^*m*^*δt*) and an additive constant, *δI*^*m*^. In our study, *m* = 1 or 2. Neuron parameters have been fitted to experimental data from 111 selected neurons according to the cell database of the Allen Brain Atlas [Allen Institute, 2018], see [Teeter et al., 2018, Billeh et al., 2020], including neuron capacity *C*, conductance *g*, resting potential *E*_L_, the length of the refractory period, as well as amplitudes *δI*^*m*^ and decay time constants *k*^*m*^ of two types of after-spike currents, *m* = 1, 2.

### 4.2 Synaptic inputs

The V1 model utilizes experimental data to specify the connection probability between neurons. The base connection probability for any pair of neurons from the 17 cell classes is provided in [Billeh et al., 2020] in a table (shown in Fig. 1B), where white cells denote unknown values. The values in this table are derived from measured frequencies of synaptic connections for neurons at maximal 75 μm horizontal inter-somatic distance. The base connection probability was then scaled by an exponentially decaying factor based on the horizontal distance between the somata of the two neurons (Fig. 1C), also derived from experimental data. The synaptic delay was spread in [1, 4] ms, as extracted from Fig. 4E of [Billeh et al., 2020] and rounded to the nearest integer as the integration step is 1 ms.

The postsynaptic current of neuron *j* was defined by the following dynamics [Billeh et al., 2020]:

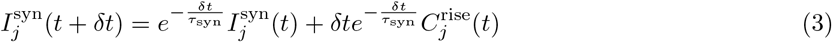

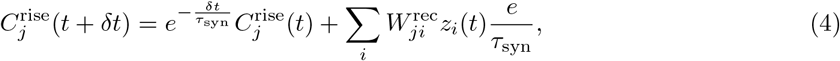

where *τ*_syn_ is the synaptic time constant, 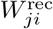 is the recurrent input connection weight from neuron *i* to *j*, and *z*_*i*_ is the spike of presynaptic neuron *i*. The *τ*_syn_ constants depend on neuron types of preand postsynaptic neurons [Billeh et al., 2020].

### 4.3 Data-driven noise model

We used a noise model that was introduced in our previous study [Chen et al., 2022]. The model was based on an empirical noise distribution that was obtained from experimental data of mice responses to 2,800 nature images [Stringer et al., 2019]. The noise currents 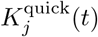 and 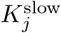 in Eq. 1 were drawn independently for all neurons from this distribution. The quick noise 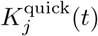 was drawn every 1 ms while the slow noise 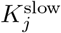 was drawn once every 600 ms. The empirical noise distribution was derived from the variability (additive noise) collected from the experimental data. A detailed mathematical analysis of this method is available in the methods and supplementary materials of [Stringer et al., 2019].

### 4.4 Projection neurons

We employed a readout population in the V1 model, whose firing activity during the response window encoded the network decisions. Each population consisted of a certain number (30 or 1) of randomly selected excitatory neurons in layer 5, located within a sphere of a radius of 55 μm (Fig. 2E).

### 4.5 Training task and injecting visual input into the model

#### LGN model

The visual stimuli were processed by a qualitative retina and LGN model, as depicted in Fig. 2C and following [Billeh et al., 2020]. Their full LGN model consists of 17,400 spatiotemporal filters that simulate the responses of LGN neurons in mice to visual stimuli [Durand et al., 2016]. Each filter generates a positive output, which represents the firing rates of a corresponding LGN neuron. We used only a subset of 2,589 of these LGN filters that provide inputs from a smaller part of the visual field to the core part of the V1 model, on which we are focusing in this study. The input images were first converted to grayscale and scaled to fit in the interval [ *−Int, Int*], where *Int >* 0. The output of the LGN model was then used as an external current input in the V1 model as follows:

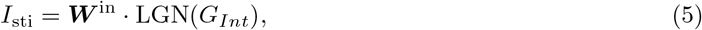

where *G*_*Int*_ represents images scaled into [*−Int, Int*] for *Int* = 2.

#### Visual-change-detection task with natural images

We designed the visual-change-detection task to be as close as possible to corresponding biological experiments while keeping them as simple as possible. In the mouse experiments of [Garrett et al., 2020b, Siegle et al., 2021], mice were trained to perform a visual-change-detection task using static natural images presented in a sequence of 250 ms with short phases (500 ms) of gray screens in between. The mice had to report whether the most recently presented image was the same as the previously presented one. To replicate this task while taking into account GPU memory limitations, we presented natural images for 100 ms each with delays between them lasting 200 ms (Fig. 2A, B). The first image was presented after 50 ms, and all images were selected from a set of 40 randomly chosen images from the ImageNet dataset [Deng et al., 2009]. The model had to report within a 50 ms time window starting 150 ms after image onset (response window) if the image had changed.

In the response window, we defined the mean firing rate of readout population as

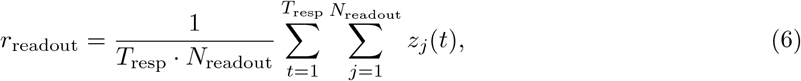

where the sum over *j* is over the *N*_readout_ = 30 projection neurons and the sum over *t* is over the time length of response window *T*_resp_ = 50 ms. If *r > r*_0_ = 0.01, the model reported a network decision that the image had changed. Otherwise, it reported no change.

#### Evidence accumulation task

A hallmark of cognitive computations in the brain is the capability to go beyond a purely reactive mode: to integrate diverse sensory cues over time, and to wait until the right moment arrives for an action. A large number of experiments in neuroscience analyze neural coding after learning such tasks (see e.g., [Morcos and Harvey, 2016a, Engelhard et al., 2019]). We considered the same task that was studied in the experiments of [Morcos and Harvey, 2016a, Engelhard et al., 2019]. There a rodent moved along a linear track in a virtual environment, where it encountered several visual cues on the left and right. Later, when it arrived at a T-junction, it had to decide whether to turn left or right. The network should report the direction from which it had previously received the majority of visual cues. To reproduce this task under the limitations of a GPU implementation, we used a shorter duration of 600 ms for each trial (Fig. S1). The right (left) cue was represented by 50 ms of cue image in which the black dots on the right (left) side of the maze. Visual cues were separated by 10 ms, represented by the gray wall of the maze. After a delay of 250 ms, the network had to decide whether more cues had been presented on the left or right, using two readout populations for left and right. The decision was indicated by the more vigorously firing readout pool (left or right) within the response window of 50 ms.

#### Injecting image without LGN

As a control measure, we also trained models in the absence of LGN. Echoing the no-LGN input approach for control model 1 in Ref. [Chen et al., 2022], we fed the image’s pixel values directly into the V1 model. This image was resized to approximate the number of active LGN channels. The pixel values were multiplied by a factor of 0.04 to align the firing activity within a suitable range.

### 4.6 Loss function

The loss function was defined as

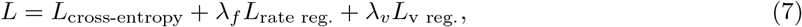

where *L*_cross-entropy_ represents the cross-entropy loss, *λ*_*f*_ and *λ*_*v*_ represent the weights of firing-rate regularization *L*_rate reg._ and voltage regularization *L*_v reg._, respectively. As an example, the cross-entropy loss of visual-change-detection task was given by

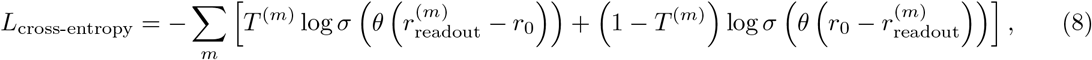

where the sum over *m* is organized into chunks of 50 ms and 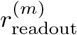 denotes the mean readout population firing rate defined in Eq. 6. Similarly, *T* ^(*m*)^ denotes the target output in time window *m*, being 1 if a change in image identity should be reported and otherwise 0. The baseline firing rate *r*_0_ was 0.01. *σ* represents the sigmoid function. *θ* is a trainable scale (*θ >* 0) of firing rate.

We used regularization terms in the loss function to penalize very high firing rates as well as values of membrane voltages that were not biologically realistic. The default values of their weights were *λ*_f_ = 0.1 and *λ*_v_ = 10^*−*5^. The rate regularization is defined via the Huber loss [Huber, 1992] between the target firing rates, *y*, calculated from the model in [Billeh et al., 2020], and the firing rates, *r*, sampled the same number of neurons from the network model:

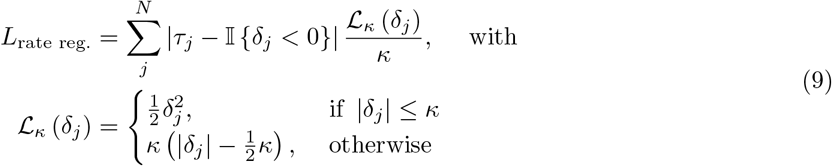

where *j* represents neuron *j, N* the number of neurons, 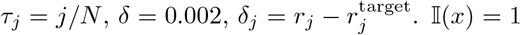 when *x* is true; I(*x*) = 0 when *x* is false.

The voltage regularization is defined through the term

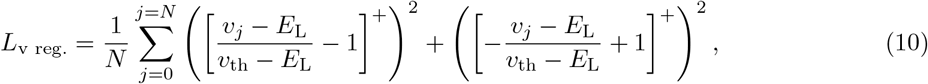

where *N* represents the total number of neurons, *v*_*j*_, the membrane potential of neuron *j*, [*· · ·* ]^+^, the rectifier function. *v*_th_ is the firing threshold of membrane potential. *E*_*L*_ the resting membrane potential.

### 4.7 Training and testing

We applied back-propagation through time (BPTT) [Chen et al., 2022] to minimize the loss function. The non-existing derivative 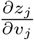 was replaced in simulations by a simple nonlinear function of the membrane potential that is called the pseudo-derivative. Outside of the refractory period, we chose a pseudoderivative of the form

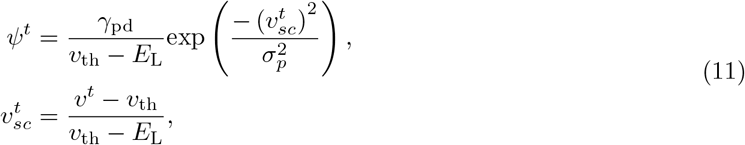

where the dampening factor *γ*_pd_ = 0.5, the Gaussian kernel width *σ*_*p*_ = 0.28. During the refractory period, the pseudo derivative was set to 0.

To demonstrate how sensitive the performance is to the scale of the surrogate derivative, we trained the model with *γ*_pd_ = 0.25 and 0.75 and kept all other hyperparameters the same. When *γ*_pd_ = 0.25, the task performance (testing accuracy) is 0.7; when *γ*_pd_ = 0.75, the testing accuracy is 0.75. Compared with the case of *γ*_pd_ = 0.5 where the task performance is 0.83, other values are worse. This demonstrates that the choice of the derivative’s scale can substantially affect gradient-based learning performance in spiking neural networks [Zenke and Vogels, 2021].

We drew a batch of visual stimuli (64) and calculated the gradient after every trial for each synaptic weight whether an increase or decrease of it (but without changing its sign) would reduce the loss function. Weights were then updated by the average gradient across the batch. This method had originally only been applied to neuron networks with differentiable neuron models and was normally referred to as stochastic gradient descent. Unless specified otherwise, initial conditions for spikes were set to zero, and membrane potentials were initialized to the resting potential. The initial configurations for **W**^in^ and **W**^rec^ were adopted from [Billeh et al., 2020]. During the training, we added the sign constraint on the weights of the neural network to keep Dale’s law. Specifically, if an excitatory weight was updated to a negative value, it would be set to 0; vice versa. In every training run, we used a different random seed in order to draw fresh noise samples from the empirical distribution, and to randomly generate/select training samples.

### 4.8 Other simulation details

The BPTT training algorithm was implemented in TensorFlow, which is optimized to run efficiently on GPUs, allowing us to take advantage of their parallel computing capabilities. We distributed the visualchange-detection task trials over batches, with each batch containing 64 trials, and performed independent simulations in parallel. Each trial lasted for 600 ms of biological time, and computing gradients for each batch took around 5 s on an NVIDIA A100 GPU. Once all batches had finished (one step), gradients were calculated and averaged to update the weights by BPTT. We define an epoch as 500 iterations/steps. This computation had to be iterated for 22 epochs to make sure the performance was saturated. This took 12 hours of wall clock time on 32 GPUs.

### 4.9 Quantifying discrepancies in sequential patterns

To assess differences between two rank orders, which signify unique network-encoded information, we leveraged Kendall’s *τ* coefficient. In the context of rank order coding, rank order essentially represents the sequence in which neurons fire. Given this, a difference in rank order between two sequences might suggest a difference in the encoded information. Kendall’s *τ* coefficient is especially apt for evaluating such ordinal data or sets with tied ranks. Kendall’s *τ* is computed as:

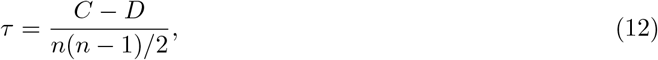

where, *C* is the count of concordant pairs, *D* signifies the number of discordant pairs, and *n* is the total observation count.

For two distinct sequences of neural firing, each rank or observation indicates the position of a neuron in the firing order. Within this framework, *x* and *y* denote two separate sequences of averaged neural activity across 200 trials. We used the times of peak activities to calculate the rank order in the network. Each observation, such as *x*_*i*_ in *x* and *y*_*i*_ in *y*, signifies the rank of a specific neuron *i* within those sequences. A pair of observations, denoted as (*x*_*i*_, *y*_*i*_) and (*x*_*j*_, *y*_*j*_), is labeled concordant if the ranks for both *x* and *y* either both increase or both decrease, which translates to the condition (*x*_*i*_ *− x*_*j*_)(*y*_*i*_ *− y*_*j*_) *>* 0. In contrast, a pair is tagged discordant if the rank of one observation decreases as the other’s increases, described by the condition (*x*_*i*_ − *x*_*j*_)(*y*_*i*_ − *y*_*j*_) *<* 0. The value of *τ* ranges from -1 to 1. A value approaching 1 signifies a strong positive association between the rank orders, a value nearing -1 indicates a strong negative association, and a value around 0 suggests no discernible association.

### 4.10 Control models

We used 7 control models. In Sec. 2.5, we introduced 3 control models.

#### Recurrent artificial neural network model (RANN)

The dynamics of a RANN can be defined as

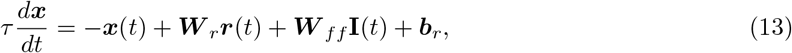

where the ***x*** is the activation of the network units and the corresponding “firing rate” is defined as

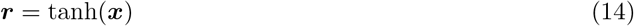

*τ* is the single-unit timescale, we used 50 ms as in [Sussillo et al., 2015, Pollock and Jazayeri, 2020]. ***W*** _*r*_ is the recurrent synaptic weights; ***W*** _*ff*_ is the feedforward synaptic weights; ***b***_*r*_ is bias. As the “firing rate” of neurons in RANN, ***r*** is in [*−*1, 1], we took the absolute value of ***r*** in Fig. S8A-C to compare with the neural activity in the V1 model. The initialization of ***W*** _*r*_, ***W*** _*ff*_ and ***b***_*r*_ are Gaussian noise *𝒩* (0, *σ*^2^). 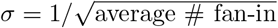 for ***W*** _*r*_ and ***W*** _*ff*_ ; *σ* = 0.1 for ***b***_*r*_. **I**(*t*) is the input to the network at time *t*. We evaluated stimulus inputs both with and without the LGN model and did not find observable differences. Unless specified otherwise, we opted for the model incorporating the LGN. The linear readout ***y***(*t*) from activities of all neurons ***r***(*t*),

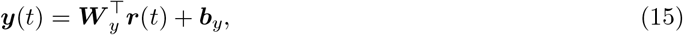

where ***b***_*y*_ is bias. ***W*** _*y*_ is the readout weight. ***W*** _*y*_ is a *N ×* 2 matrix; *N* is the number of recurrent neurons; 2 is the number of possible decisions. The number of neurons and synapses in our model are the same as those in Billeh’s model [Billeh et al., 2020]. We randomly shuffled the connectivity or kept it the same as in Billeh’s model.

The loss function was defined as

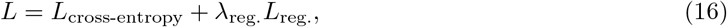

where *L*_cross-entropy_ represents the cross-entropy loss which was defined in Eq. 8, *λ*_reg._ represents the weight of activation regularization *L*_reg._. The activation regularization was defined as

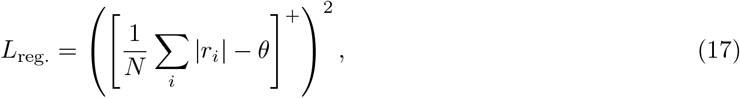

where *r*_*i*_ is the “firing rate” of neuron *i. N* is the number of neurons and *θ* is a threshold (*θ* = 0.01), unless otherwise stated. [*· · ·* ]^+^ is the rectifier linear unit function. The value of *θ* was determined as 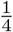 of the mean value of |*r*| when training the RANN model without regularization. We used the same training methods as in the V1 model but the learning rate is 10^*−*4^. We also used *dt* = 5 ms to alleviate the vanishing gradient problem, as the one used in [Sussillo et al., 2015].

#### Recurrent convolution neural network (RCNN)

We used RCNN [Wang and Hu, 2021] as another control model, the same one used in [Chen et al., 2022], inspired by abundant recurrent connections in the visual systems of animals. The gates control the amount of context information inputted to the neurons. We used the code, GRCNN-55 (weight sharing). The number of neurons in RCNN is comparable to the V1 model [Billeh et al., 2020]. The loss function only contains the cross entropy term in Eq. 8, without any regularization. We used the same training methods as in the V1 model but the learning rate was 1 *×* 10^*−*6^. The batch size was 64; the number of training epochs was 100. We also used *dt* = 5 ms to alleviate the vanishing gradient problem, as the one used in [Sussillo et al., 2015]. We considered the neuronal activation after the activation function (ReLu) as the “firing rate” of neurons in RCNN. We evaluated stimulus inputs both with and without the LGN model and did not find observable differences. Unless specified otherwise, we opted for the model incorporating the LGN.

#### Generic recurrent spiking neural network (RSNN)

We removed the laminar spatial structure with distance-dependent connection probabilities of the V1 model and replaced them with an equal number of randomly chosen connections. This model is a randomly connected recurrent network of standard LIF neurons with the same number of neurons and connections as the V1 model. The membrane time constant of these LIF neurons is 10 ms, and their refractory period is 5 ms long. These are common values for spiking neural network models. We evaluated stimulus inputs both with and without the LGN model and did not find observable differences. Unless stated otherwise, we opted for the model incorporating the LGN.

In Sec. 2.6, we used 5 control models which were also used in [Chen et al., 2022]. In control model 1, we removed the LGN model, and directly injected pixel values of the image into the V1 model. The image was resized to a number of pixels that roughly matched the number of LGN channels (2544). The pixel values were scaled by a factor 0.04 to bring the resulting firing activity into a reasonable regime. In control model 2, we removed the diversity of the 111 data-based neuron types in the V1 model, replacing them with one generic model for excitatory neurons (the excitatory neuron on L2/3, node type id in Allen brain atlas: 487661754) and one for inhibitory neurons (PV neuron on L2/3, node type id: 484635029). In control model 3, we removed instead the laminar spatial structure with distance-dependent connection probabilities of the V1 model, replacing them with an equal number of randomly chosen connections. Control model 4 is the aforementioned RSNN. Control model 5 is a variation of this RSNN where excitatory and inhibitory neurons are distinguished like in the V1 model, and Dale’s law is observed during training. This type of model is sometimes seen as an intermediate step from generic RSNNs toward more biologically oriented network models. The detailed settings are shown in Table S1. All control models were trained in the same way as the V1 model, i.e., for the same tasks and loss function, including the same sparsity regularization term, and using the same hyperparameters for training.

### 4.11 Demixed principal component analysis

Demixed principal component analysis (Demixed PCA) is a statistical method that decomposes highdimensional neural data into a set of orthogonal latent variables, each of which captures a unique aspect of the neural response [Kobak et al., 2016]. Briefly, let **X***n × T* be the neural data matrix, where *n* is the number of neurons and *T* is the number of time points. Let **S***n × C × T* be the tensor of stimulus conditions, where *C* is the number of experimental conditions. The goal of Demixed PCA is to find a low-dimensional latent space **Y**_*d×T*_, where *d* is the number of latent variables, that captures the majority of the variance in the neural data **X**, while also separating out the variance that is specific to each experimental condition in **S**.

In Fig. 3D, the principal component used for the projection arises by analysis of the first eigenvector of the covariance matrix that reflects variation through joint dependencies of the network decision and relative timing, see marginalization procedure of [Kobak et al., 2016]. Hence, this matrix does not reflect variation that is caused only through the course of time within a trial or through the network decision alone. Moreover, to emphasize the formation of the network decision, we include in the computation of the aforementioned covariance matrix only data within [-50, 50] ms of the image presentation.

### 4.12 Mutual information

To estimate the mutual information between single neuron activity and the network decision, we binned the spike counts of each neuron into 10 uniformly distributed bins between the minimum and maximum spike count observed for that neuron within 50 ms windows. We then established an empirical joint distribution for the binned spike count and the network decision and computed the mutual information using the below formula.

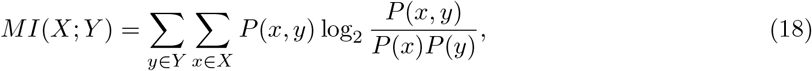

where *X* is the set of firing activities of each neuron within a 50 ms window, and *Y* is the set of network decisions (either change or no-change). *P* (*x, y*) is the joint probability distribution of spike count and network decision, while *P* (*x*) and *P* (*y*) are the marginal probability distributions of spike count and network decision, respectively.

To estimate *P* (*x, y*), we calculated the spike count of each neuron within a 50 ms window and established an empirical joint distribution by counting the number of occurrences of each possible combination of spike counts and network decisions across 100,000 trials. We then normalized the joint distribution to obtain a probability distribution. We estimated *P* (*x*) and *P* (*y*) in a similar way by counting the number of occurrences of each possible value of spike count and network decision, respectively, across all trials.

### 4.13 UMAP

We applied an exponential filter with a time constant of 20 ms to the spike output of each neuron for 8 new images that had not been used during training. We then discarded all but the 1,500 most important principal components of these network states (explain 38% of variance on visual-change-detection task and 41% of variance on evidence accumulation task; compromise to the memory consumption), and embedded these into 2D space by UMAP. These projected network states were recorded for every ms, represented by a dot in Fig. 5.

UMAP (Uniform Manifold Approximation and Projection) is a nonlinear dimensionality reduction algorithm designed to maintain the inherent structure of high-dimensional data when projected into a lower-dimensional space [McInnes et al., 2018]. Unlike linear techniques, UMAP focuses on preserving the pairwise distances between neighboring data points. In other words, it strives to conserve the local relationships or similarities among points in the original high-dimensional data. In this study, UMAP was used to embed the network activity during task performance into 2D space. Specifically, we used the UMAP implementation from the Python library umap, with the following parameters: n neighbors = 200, min dist = 0.1, and metric = euclidean.

The resulting 2D embeddings represent the low-dimensional trajectories of the network states during image processing, and were recorded for every ms. Each point in the 2D space represents a network state at a given time point, and is displayed as a dot in Fig. 5. The trajectory of the network states can be visualized by connecting these dots in chronological order.

### 4.14 Additional information for figures

#### Normalized activity in Figs. 3, S8, S9, S12 and S11

Spiking activity at a specific relative time step, regarding image presentation, was averaged over 200 trials. These average activities per time step were then normalized with the maximum values of their average activation.

#### Fig. 6B and C

7 neurons shown in Fig. 6B and C were selected from the 20 early-informer-neurons with the largest MI that represented each of the 4 neuron classes, taking within each neuron class (excitatory, PV, Htra3, or Sst neurons) the ones with the largest MI.

## Acknowledgements

We would like to thank Anton Arkhipov, Shinya Ito, and Omid Zobeiri for useful discussions on reanalyzing neuropixel data from Allen brain institute and thank Yuqing Zhu for helpful discussions. We also thank Sandra Diaz for advice and help in using supercomputers. This research was partially supported by the Human Brain Project (Grant Agreement number 785907) of the European Union and a grant from Intel. Computations were carried out on the Human Brain Project PCP Pilot Systems at the J·lich Supercomputing Centre, which received co-funding from the European Union (Grant Agreement number 604102).

## Supplementary Information for

### Supplementary Note 1: Details to the evidence accumulation task

Besides the visual-change-detection task, we trained another task in the V1 model and control models: evidence accumulation task (Methods) [Morcos and Harvey, 2016a, Engelhard et al., 2019]. Much like the visual-change-detection task in the V1 model (Fig. 3), the evidence accumulation task utilizes the temporal sequence of peak activity to perform computations (Fig. S6A-C). Note that in Fig. S6A-C, we used the sequence of left-right-left-left-left for the turning-left trials and the sequence of left-right-leftright-right for the turning-right trials, in order to better demonstrate the difference between two rank orders. The sequential patterns for turning left and turning right are distinctly different, as evidenced by Kendall’s *τ* coefficient being close to 0 (Table 2).

Using a similar method to calculate the mutual information between neural activity and network decision, we discovered that silencing a relatively small number of neurons (for instance, 100 with high mutual information: early-informer-neurons) dramatically reduces task performance (Fig. S6D). We selected neurons to silence based on the descending order of their mutual information with the network decision calculated within a specific time period. We observed that neuronal activities carry increasingly higher information as they approach the response window, reflected by the severe reduction of task performance (Fig. S6D). Also, we computed the mutual information between the neural activity over a temporal segment and the current state of accumulated cues (i.e., when the number of left cues is greater than, less than, or equal to the number of right cues). We found a similar effect when silencing neurons that exhibited high values of this type of mutual information.

To illustrate the computational progression of the V1 model in the evidence accumulation task, we employed UMAP to embed its activity vectors into two dimensions, mirroring the method used in the visual-change-detection task. For ease of visualization and analysis, we held the first three cues constant as left-right-left. The processing of each unique image generates two bundles of trajectories, with trial-to-trial variations arising from the sequence of cues and intrinsic network noise (Fig. S14A). These trajectories exhibit three nested bifurcations, signifying the computational progression of the dynamical system (see Fig. S14B). By silencing 100 early-informer-neurons (neurons with the highest mutual information with the network decision in a certain window), these bifurcations can be flipped (Fig. S14C). This behavior mirrors that observed in the visual-change-detection task (Fig. 5).

We also trained the control models in Sec. 2.5 and 2.6 for evidence accumulation task. As demonstrated in Table 2, the quantitative measures suggest the different computational organization in control models from the mouse brain and the V1 model.

### Supplementary Note 2: Details to control models for the visual-change-detection task

Supplement for Sec. 2.5, we show here that RCNN and RSNN are not able to reproduce the two previously discussed fingerprints of cortical computations: a short period of peak activity for most neurons, a characteristic sequential order of this peak activity according to the trial type, and the functional segregation.

#### RCNN

During both change and no-change conditions, the activity patterns of the RCNN, as depicted in Fig. S11, are notably non-sparse. The pronounced similarity between these conditions complicates the differentiation based on rank order. This observation aligns with the elevated Kendall’s *τ* values presented in Table 1.

#### Generic RSNN

As a control model, we removed the laminar spatial structure with distance-dependent connection probabilities of the V1 model and replaced them with an equal number of randomly chosen connections. This model is a randomly connected recurrent network of standard LIF neurons with the same number of neurons and connections as the V1 model. The membrane time constant of these LIF neurons is 10 ms, and their refractory period is 5 ms long. These are common values for spiking neural network models. As shown in Fig. S12A-C, our findings differ significantly from the V1 model presented in Fig. 3A-C. A large number of neurons remain inactive during network computation. Using the analysis methodology from [Driscoll et al., 2017, Koay et al., 2022], the trial type’s influence on the sequence of peak neuronal activity is weaker than in the V1 model (Fig. 3A-C). The rank order similarity for both change and no-change conditions, as indicated by Kendall’s *τ* coefficient, is notably higher (Table 1), suggesting the distinct computational organization in RSNN. In addition, we estimated mutual information same as in the V1 model and gradually silenced early-informer-neurons with the highest MI within the [50, 100] ms interval. Fig.S12D and the functional segregation index (Table1) show that the accuracy decreases, though not as significantly as in the V1 model.

**Table S1:** The control model settings.

| Model name | Neuron model | Connectivity<sup>a</sup> | Neuron types<sup>b</sup> | Weight sign constraint | Input via LGN model |
| --- | --- | --- | --- | --- | --- |
| V1 model | GLIF<sub>3</sub> | V1 | V1 | Yes | Yes |
| V1 model without LGN (control model 1) | GLIF<sub>3</sub> | V1 | V1 | Yes | No |
| V1 model without neuron diversity (control model 2) | GLIF<sub>3</sub> | V1 | 1E 1I | Yes | Yes |
| V1 model without laminar structure (control model 3) | GLIF<sub>3</sub> | random | V1 | Yes | Yes |
| RSNN (control model 4) | LIF<sup>c</sup> | random | single | No | Yes |
| RSNN with E/I distinction (control model 5) | LIF | random | single | Yes | Yes |

(a) V1: the connectivity in the V1 model; random: randomly selected connectivity but the number of synaptic connectivity is the same as in the V1 model.
(b) V1: diverse neuron types as in the V1 model (in total 111 types); 1E 1I: all excitatory neurons are the same (an excitatory neuron on L2/3, neuron id in Allen brain atlas: 487661754) and all inhibitory neurons are the same (PV neuron on L2/3, neuron id:484635029); Single: all neurons are the same.
(c) LIF represents standard LIF neuron model.

**Figure S1:**
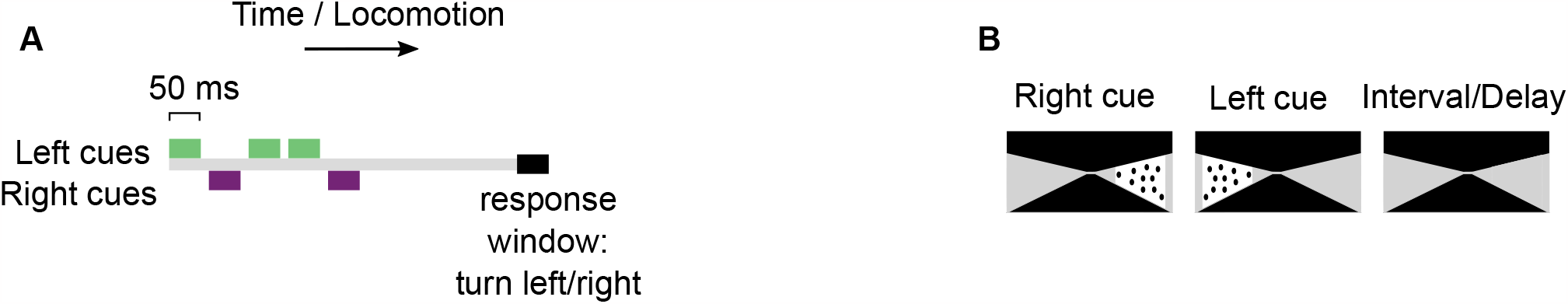
Schematic diagram of evidence accumulation task. **(A)** In the evidence accumulation task, 5 cues were presented sequentially, and after a delay, a corresponding readout pool had to indicate through stronger firing whether the majority of the cues had been presented on the left or the right. **(B)** Cue and delay images simulate a mouse navigating through a T-maze.

**Figure S2:**
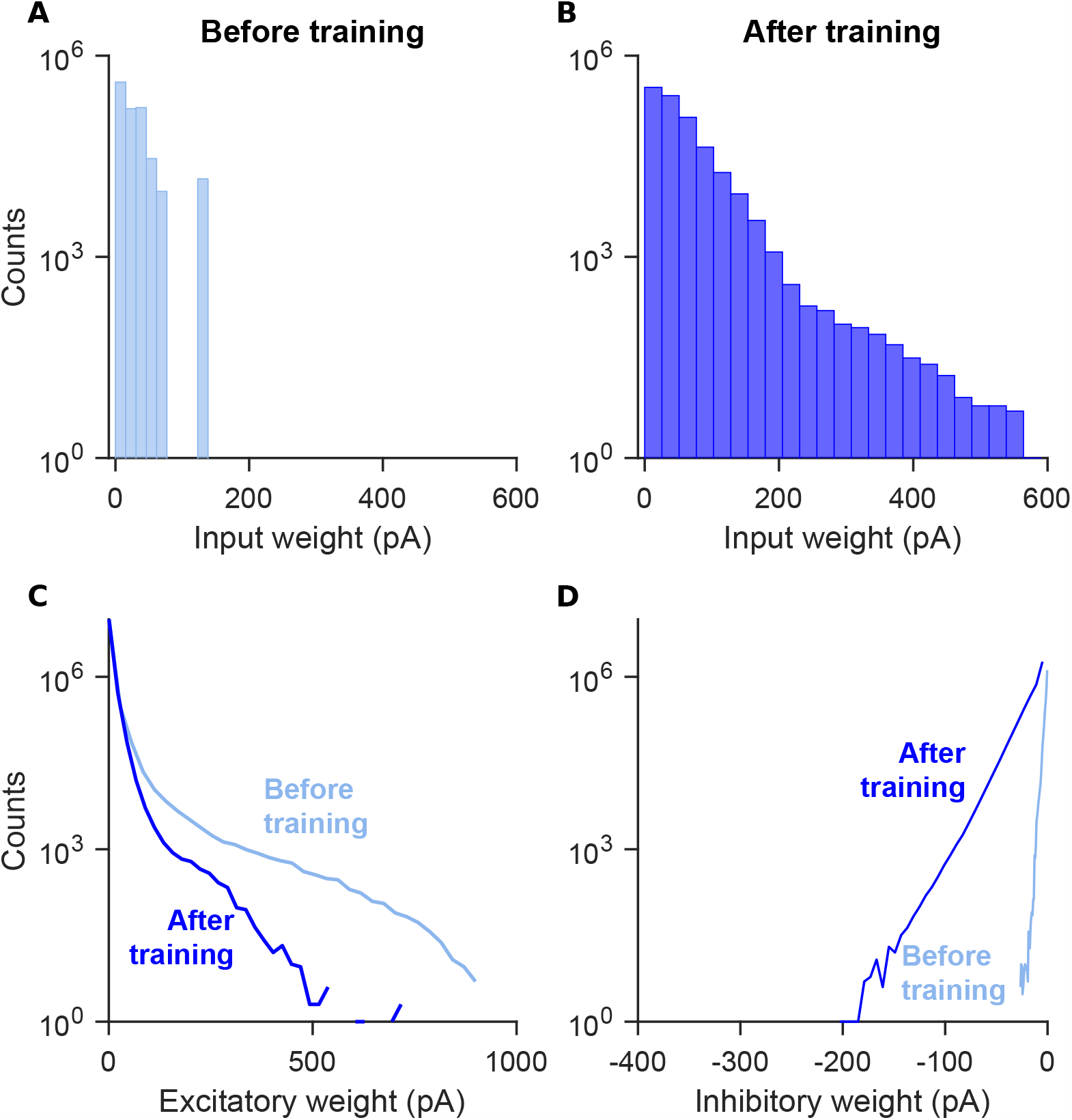
Changes in the distribution of synaptic weights through training. **(A)** Distributions of input weights before training. **(B)** Same as **(A)**, but after training. The mean of input weights increases from 24.1 to 25.2 through training. **(C)** Distributions of excitatory weights in V1 model before and after training. **(D)** Distributions of inhibitory weights in V1 model before and after training. Note that the weights before training were given by [Billeh et al., 2020].

**Figure S3:**
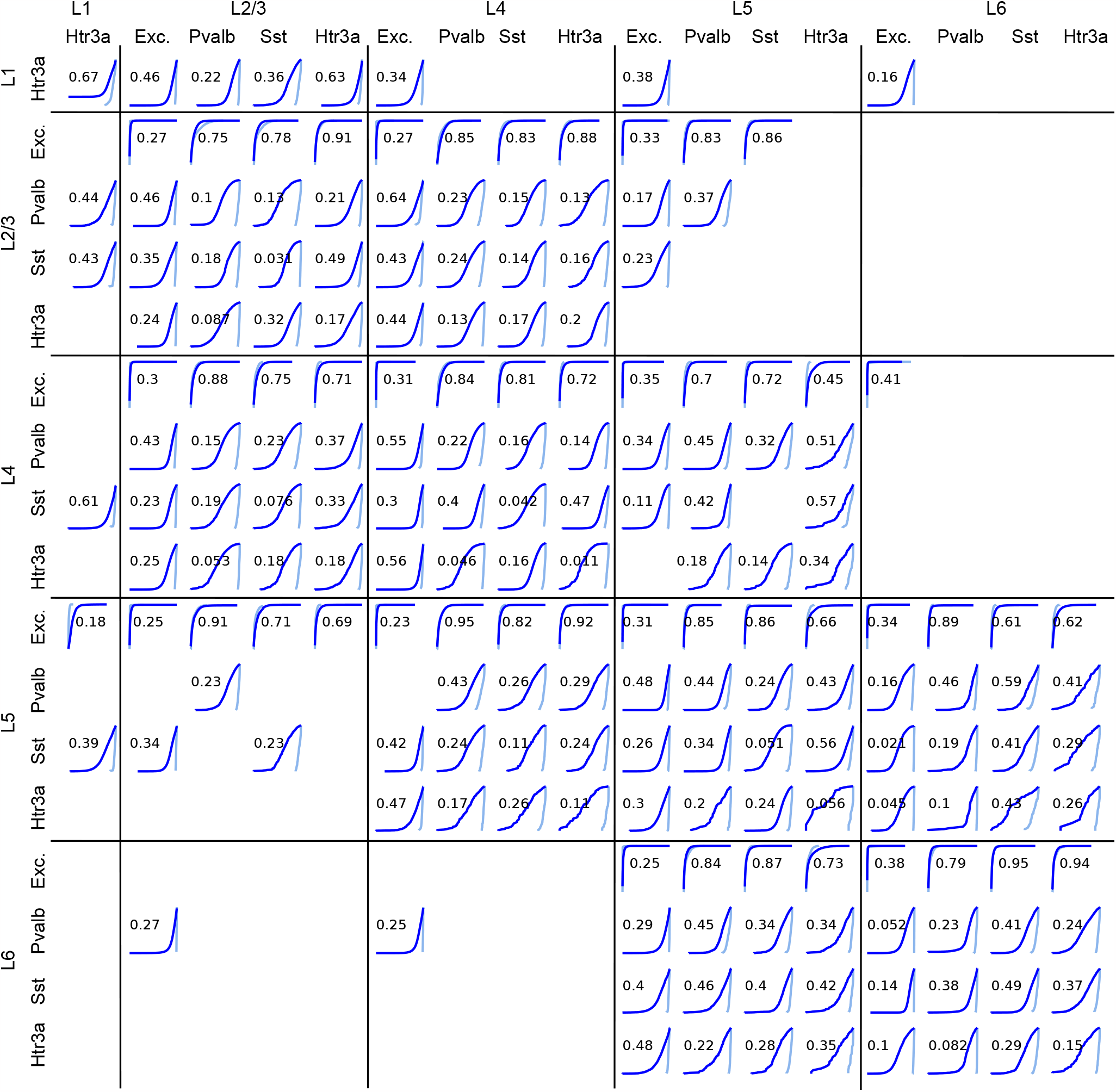
Distribution of recurrent synaptic weights between each pair of populations before (light blue) and after learning (dark blue). Each row represents a presynaptic neuron population, and each column represents a postsynaptic neuron population. The histogram represents the distribution of synaptic weights of all synaptic connections that share the same presynaptic and postsynaptic neuron population. Vertical axis in each panel is log-scale. Horizontal axis is linear scale and horizontal range is from the smallest value to the largest value of each population. The number is 1 *− D* where *D* is from the Kolmogorov–Smirnov test, quantifying the similarity between distributions [Billeh et al., 2020]. Exc., excitatory neurons.

**Figure S4:**
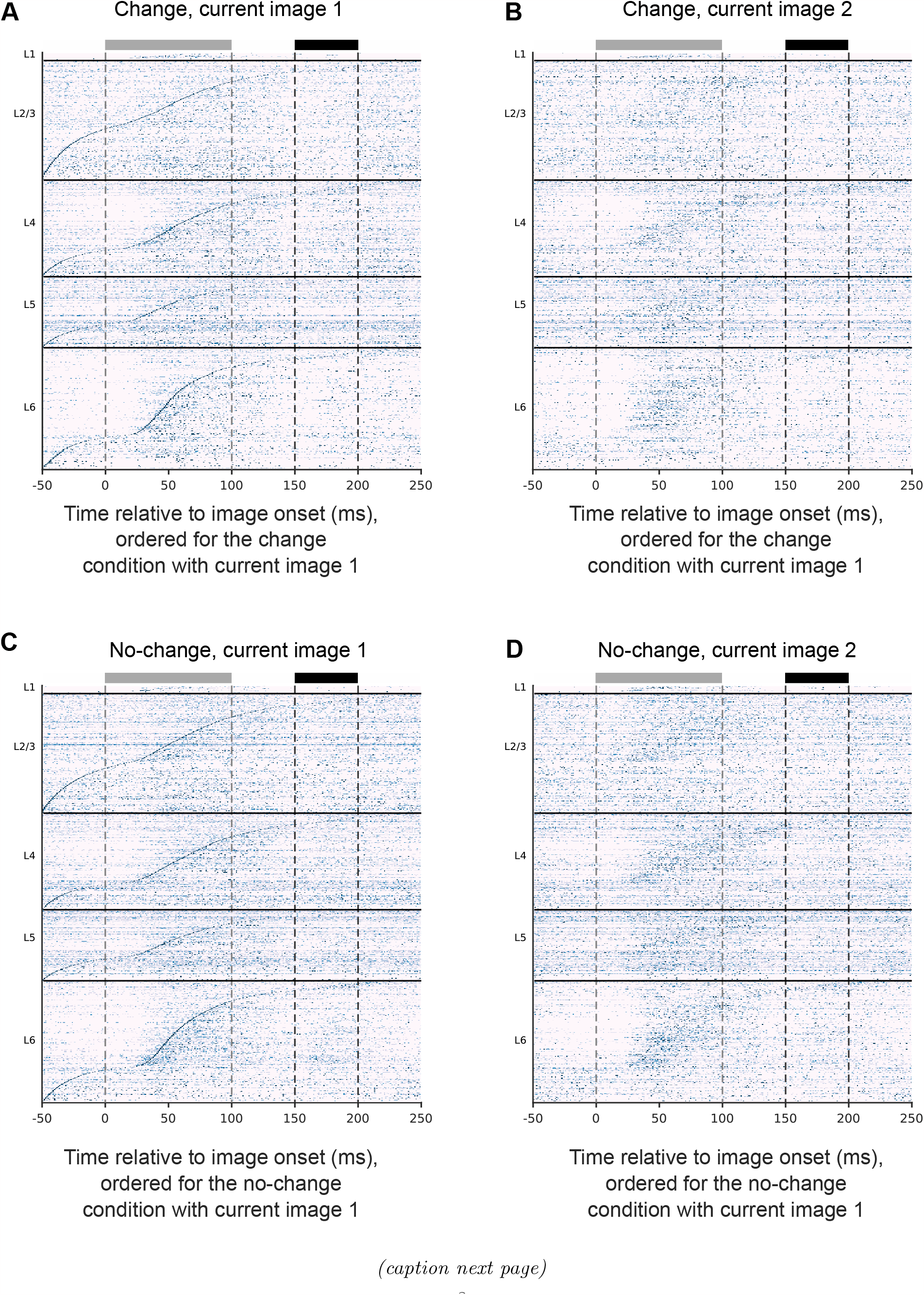
The temporal order of peak activity encodes in the V1 model also information about the identity of the current image in the visual-change-detection task. **(A)** Normalized average responses over 200 trials with the change condition and the same current image (image 1) but different preceding images, with neurons ordered according to the time of their peak activity. The gray and black bars at the top denote the image presentation and response windows, respectively. **(B)** Same as in **(A)**, but all trials have the same current image 2 and neurons were ordered as in **(A)**. The resulting blurred sequence indicates that the order of peak activity of neurons is different for images 1 and 2, also within the same trial type (change condition). **(C)** and **(D)** Same as in **(A)** and **(B)**, respectively, but for the no-change condition.

**Figure S5:**
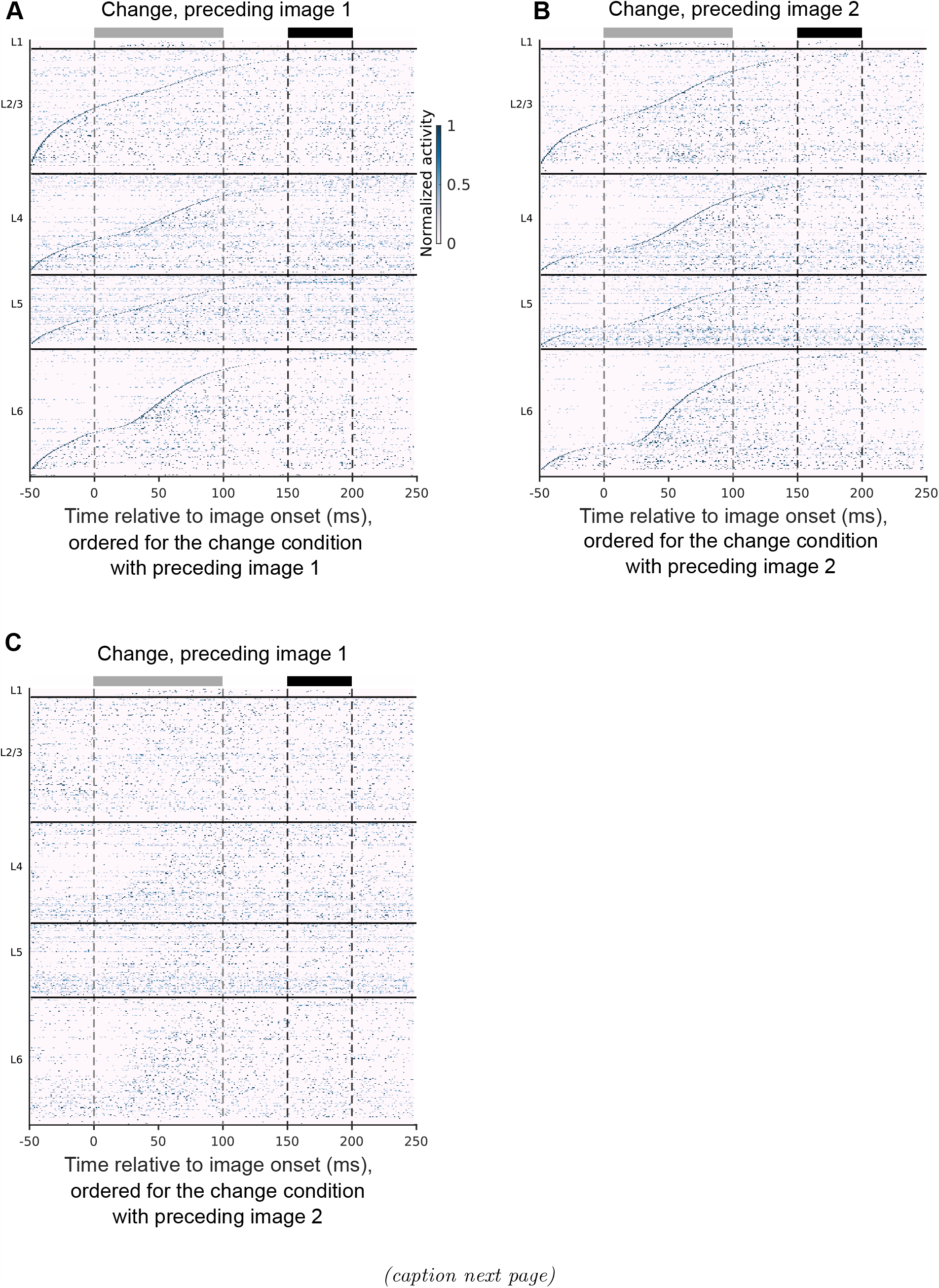
The temporal order of peak activity encodes in the V1 model also information about the identity of the preceding image in the visual-change-detection task. We have shown in Fig. 3 that the temporal order of peak activity in the V1 model has characteristic differences for different trial types (change or no-change). We show here that this temporal or rank-order coding is even more refined: the order contains in addition information about the identity of the preceding image. As preceding images were varied, we only show the change condition. **(A)** Normalized average responses over 200 trials with same preceding image (image 1) but different current images, with neurons ordered according to the time of their peak activity. The gray and black bars at the top denote the image presentation and response windows, respectively. **(B)** Same as in **(A)**, but all trials have the same preceding image 2, with neurons ordered according to the time of their peak activity for image 2. **(C)** Same data as in **(A)** but with neurons ordered as in **(B)**.

**Figure S6:**
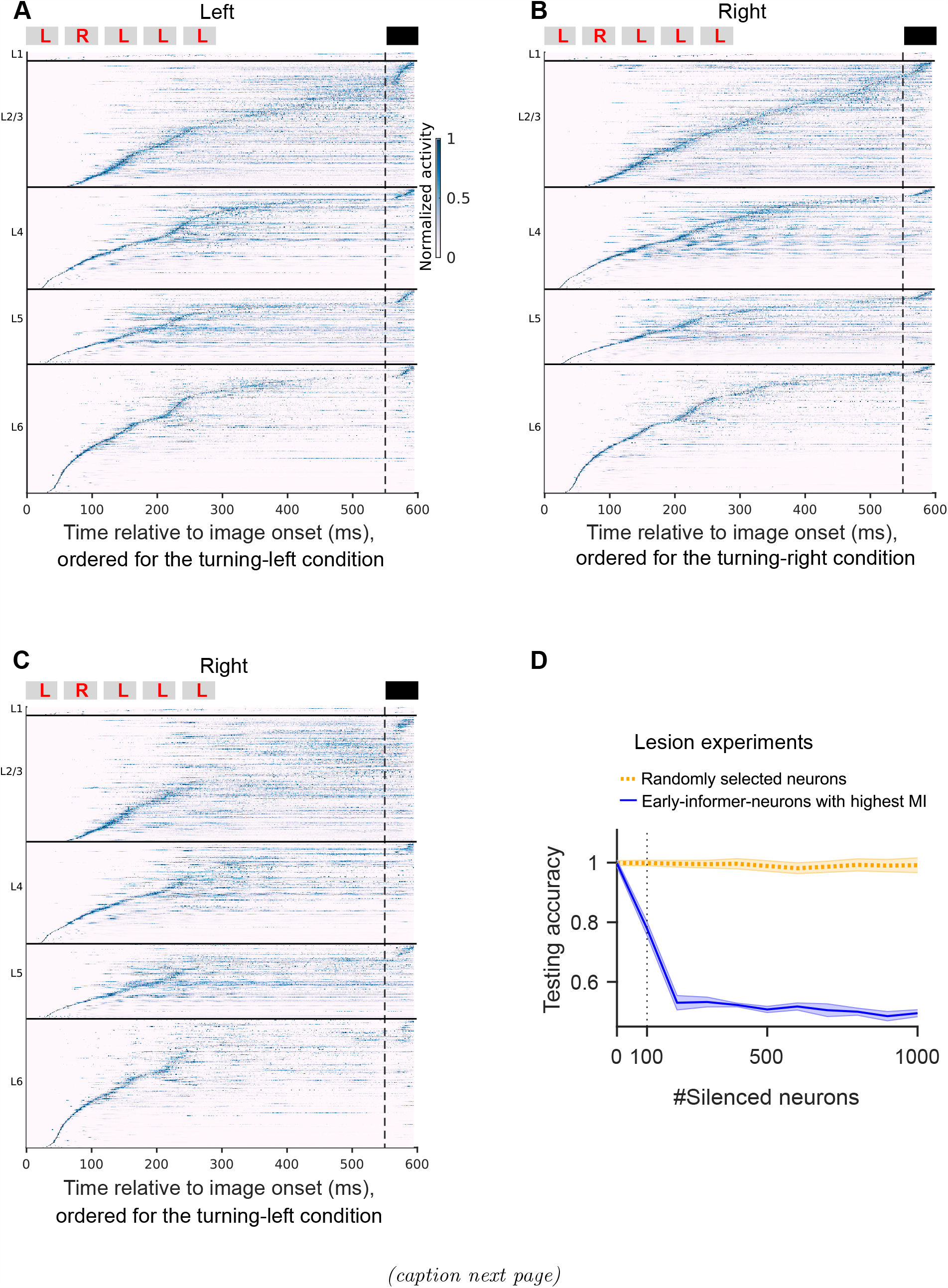
Temporal organization of computations of evidence accumulation task in the V1 model. **(A)** Normalized average responses over 200 trials with the turning-left condition, with neurons ordered according to the time of their peak activity under the turning-left condition. The gray and black bars at the top denote the cue presentations and response windows, respectively. The red letters, L (left) / R (right) in gray bars represent the meaning of cue. To reduce the variability, we used the sequence, LRLLL for the turning-left trials. **(B)** Same as in **(A)**, but for the turning-right condition, with neurons ordered according to the time of their peak activity for the turning-right condition. To reduce the variability, we used the sequence, LRLRR for the turning-right trials. **(C)** The same data as in **(B)** for the turning-right condition, but with neurons ordered as in **(A)**. The resulting blurred sequence indicates that the order of peak activity of neurons is quite different for the turning-left and turning-right conditions. **(D)** Causal impact of specific neurons on the network decision: Task performance quickly decreases when early-informer-neurons, ranked by their MI during the third cue presentation coinciding with the network decision, are silenced (represented by the blue curve). On the other hand, task performance is robust to silencing the same number of randomly selected neurons (dotted yellow curve). Both curves show average values for 10 V1 models where different random seeds were used. The shaded area represents the SEM across 10 models.

**Figure S7:**
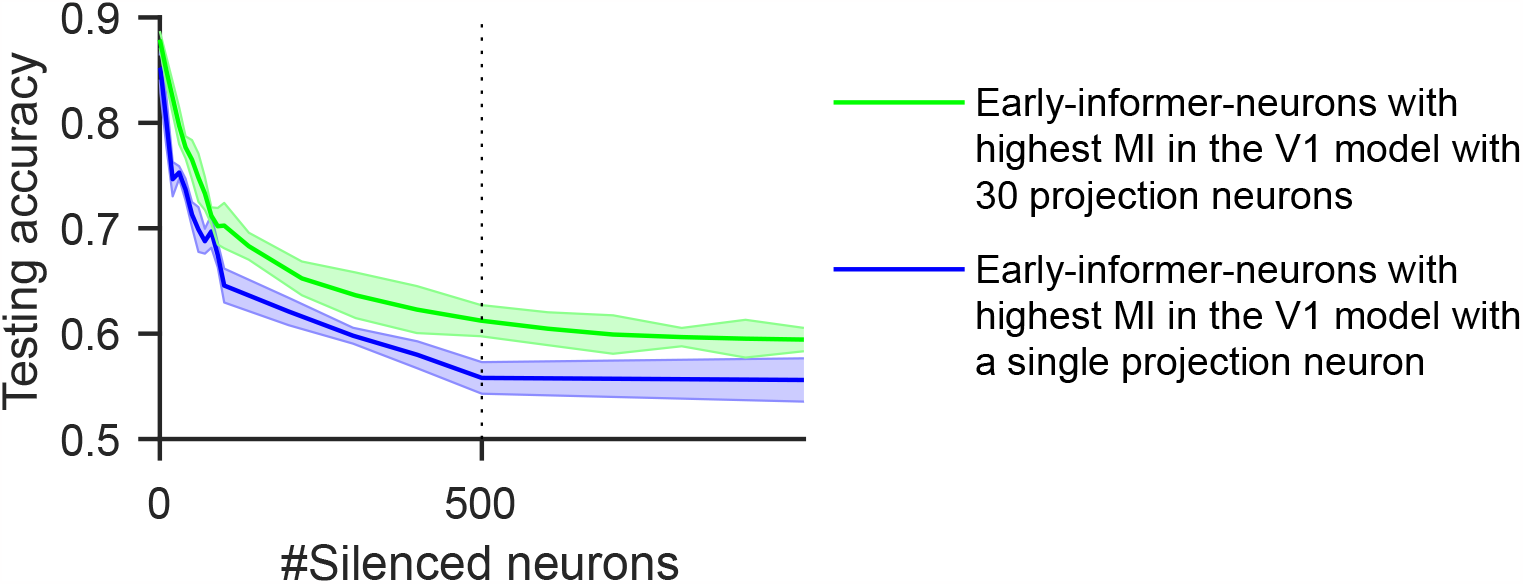
Functional segregation experiments for two versions of the trained V1 model with 30 and 1 projection neurons in the visual-change-detection task. Task performance quickly decreases when early-informer-neurons are silenced (in the descending order of their MI with the network decision) in both versions. The 30-readout-neuron model has also after silencing a given number of neurons slightly higher accuracy on test images than the single-readout-neuron model, consistent with Fig. 2F. Both curves show average values for 10 V1 models where different sets of training data were used. The shaded area represents the SEM across 10 models trained with 10 different datasets.

**Figure S8:**
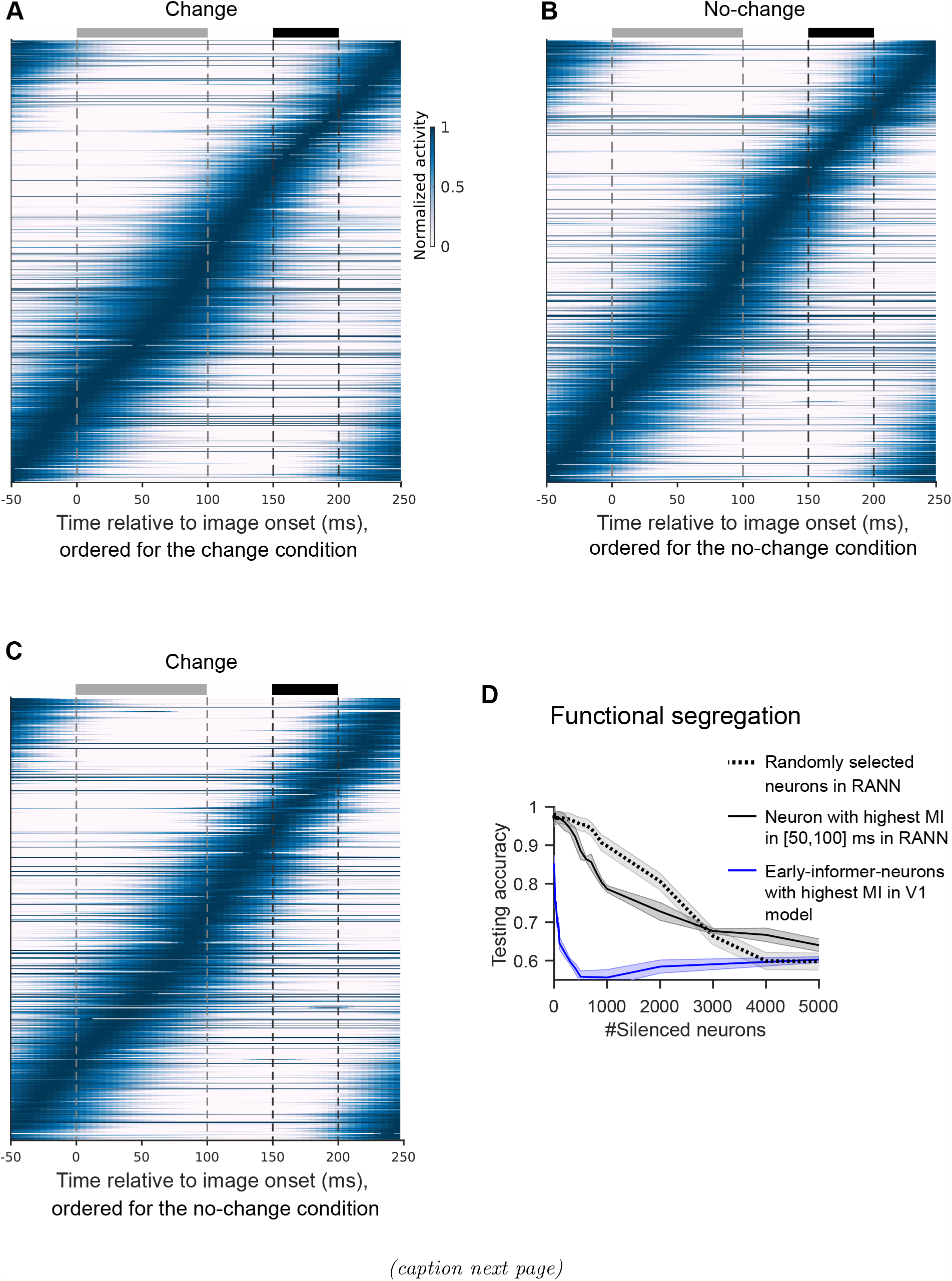
Temporal organization of the visual-change-detection task in the RANN model. **(A)** Neuronal activity in the equally large RANN model was plotted for the same network inputs and task condition as in Fig. 3A. Here, the activation regularization was not used in the RANN; in other words, the weight of activation regularization is 0 (Methods). **(B)** Same as in **(A)**, but for the no-change condition as in Fig. 3B. **(C)** Same data as in panel **(A)**, but with neurons ordered as in panel **(B)**. In contrast to Fig. 3, little difference emerges between panels **(A)** and **(C)**, indicating that the order of peak activity is less dependent on the task condition in the RANN. **(D)** Lesion experiments corresponding to those in the V1 model (Fig. 3E). The blue curve for the V1 model is the same as in Fig. 3E. One clearly sees that the network decision is substantially less sensitive to the activity of 100 neurons. The shaded areas represent SEM across 10 RANNs where different sets of training data are used.

**Figure S9:**
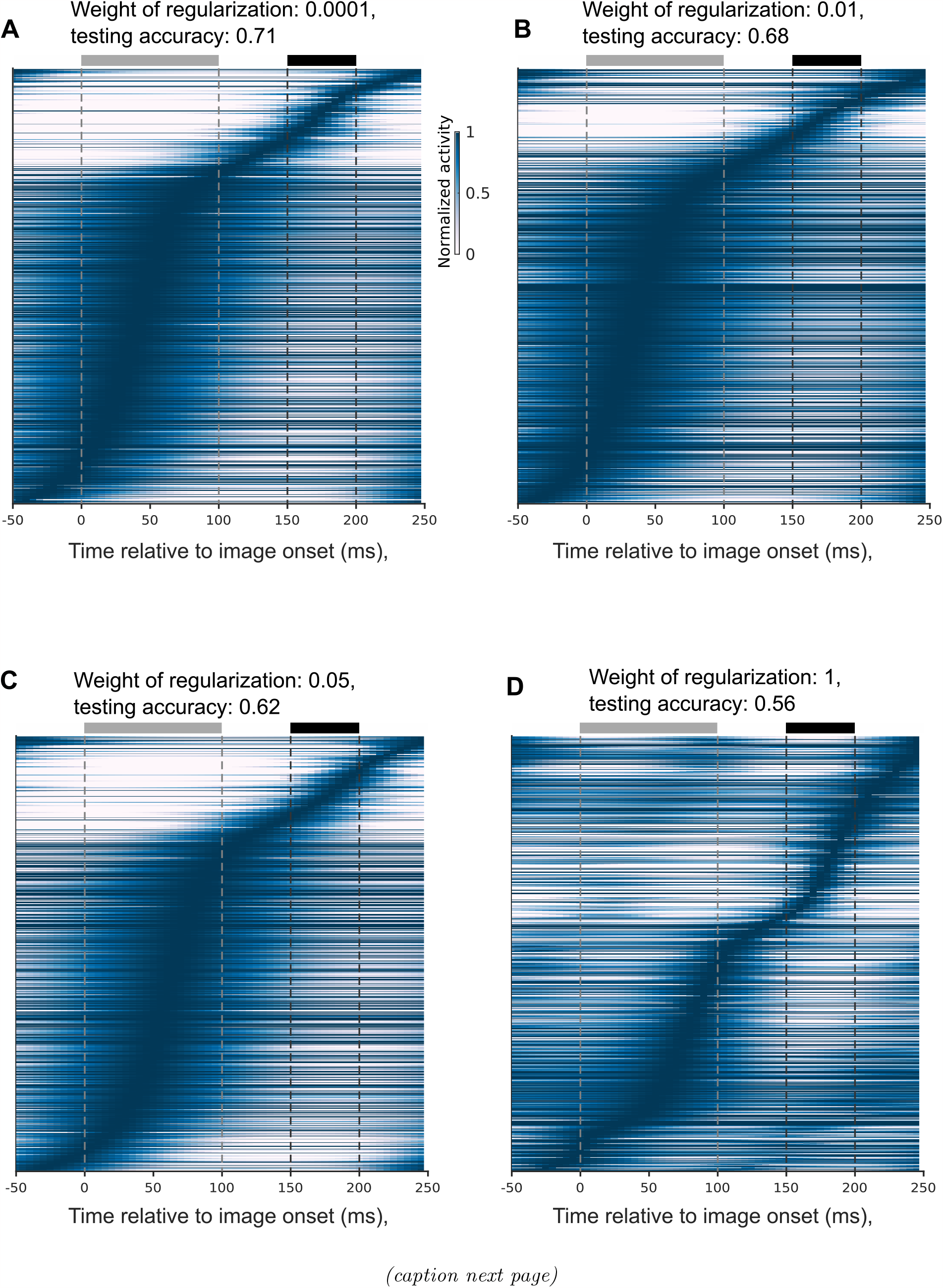
Different weights of the regularization term in RANN cannot produce sparse sequential neural activity while maintaining high task performance in the visual-change-detection task. **(A-D)** The normalized average responses of 200 change-condition trials in the RANN with different weights of regularization are plotted over time, with neurons ordered based on the time of their peak activities under the change condition. One sees that training the RANN for the same task with different weights of the regularization term (Methods) does not produce a sparse sequential neural activity as in the experimental data and the V1 model. Furthermore, more aggressive regularization strongly reduces task performance.

**Figure S10:**
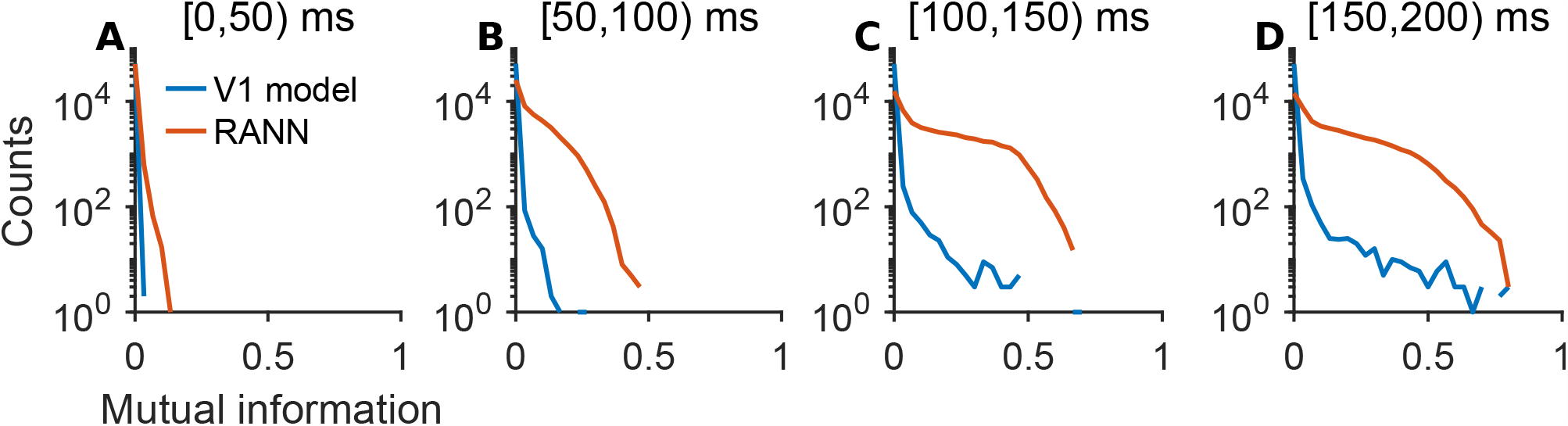
Comparison of mutual-information distributions among neurons in the V1 Model and the RANN performing visual-change-detection task. **(A-D)** The number of neurons that have a given level of MI with the network output is shown for each 50-ms window in the V1 model and the RANN (image onset was at time 0). One clearly sees that for each of these time windows that are by 1 or 2 orders of magnitude more neurons that have high MI with the network output. This explains also why so much more neurons need to be silenced in the RANN in order to reduce the task performance of the network to a given level.

**Figure S11:**
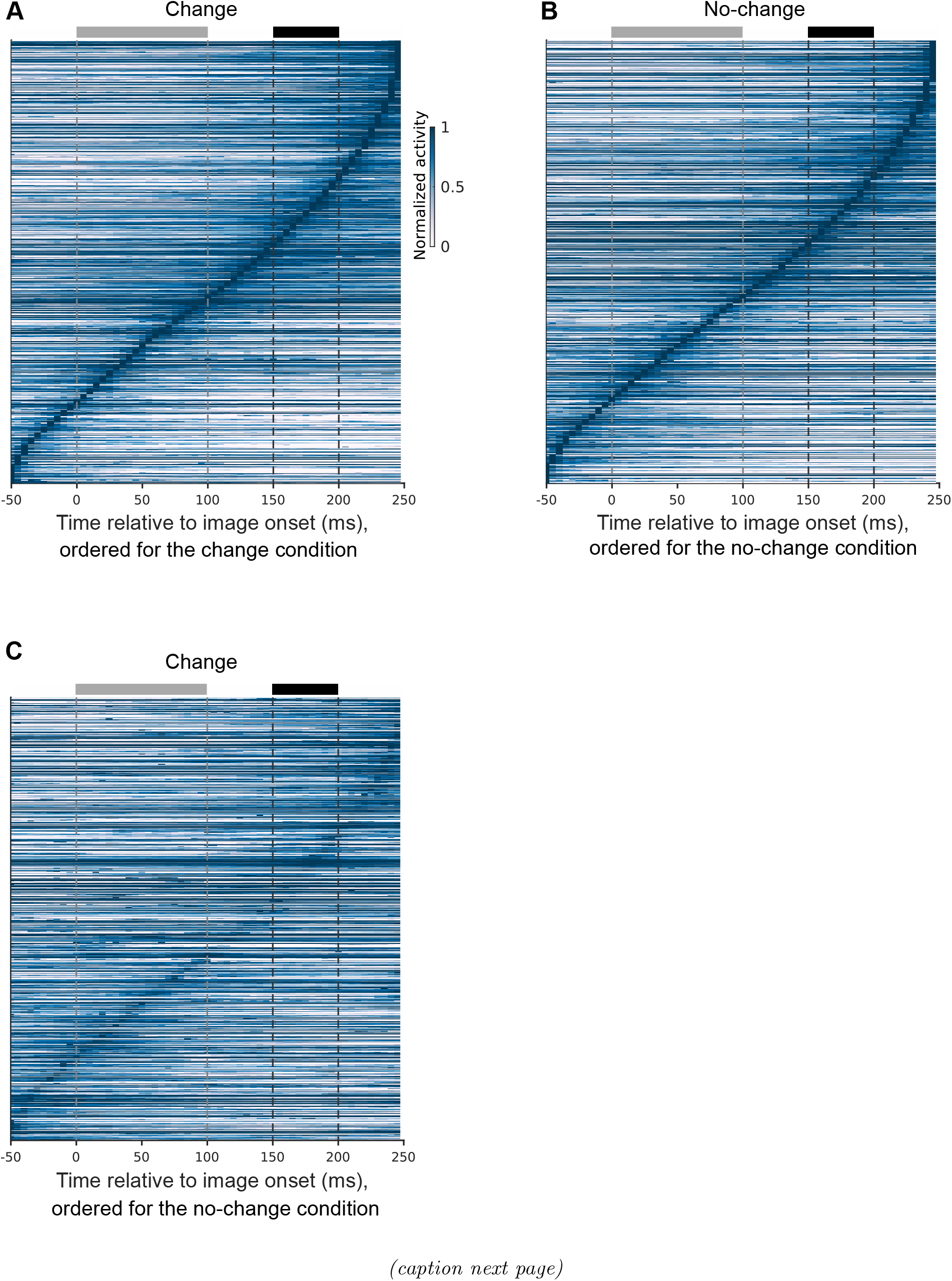
Temporal organization of computations in recurrent convolution neural network (RCNN) on visual-change-detection task. **(A)** Normalized average responses of 200 change-condition trials in RCNN are plotted over time, with neurons ordered based on the time of their peak activities under change condition. The gray and black rectangles denote the image presentation and response windows, respectively. **(B)** Same as in **(A)**, but for the no-change condition. **(C)** Normalized neural activities in the change condition are visualized with the order of neurons in the no-change condition.

**Figure S12:**
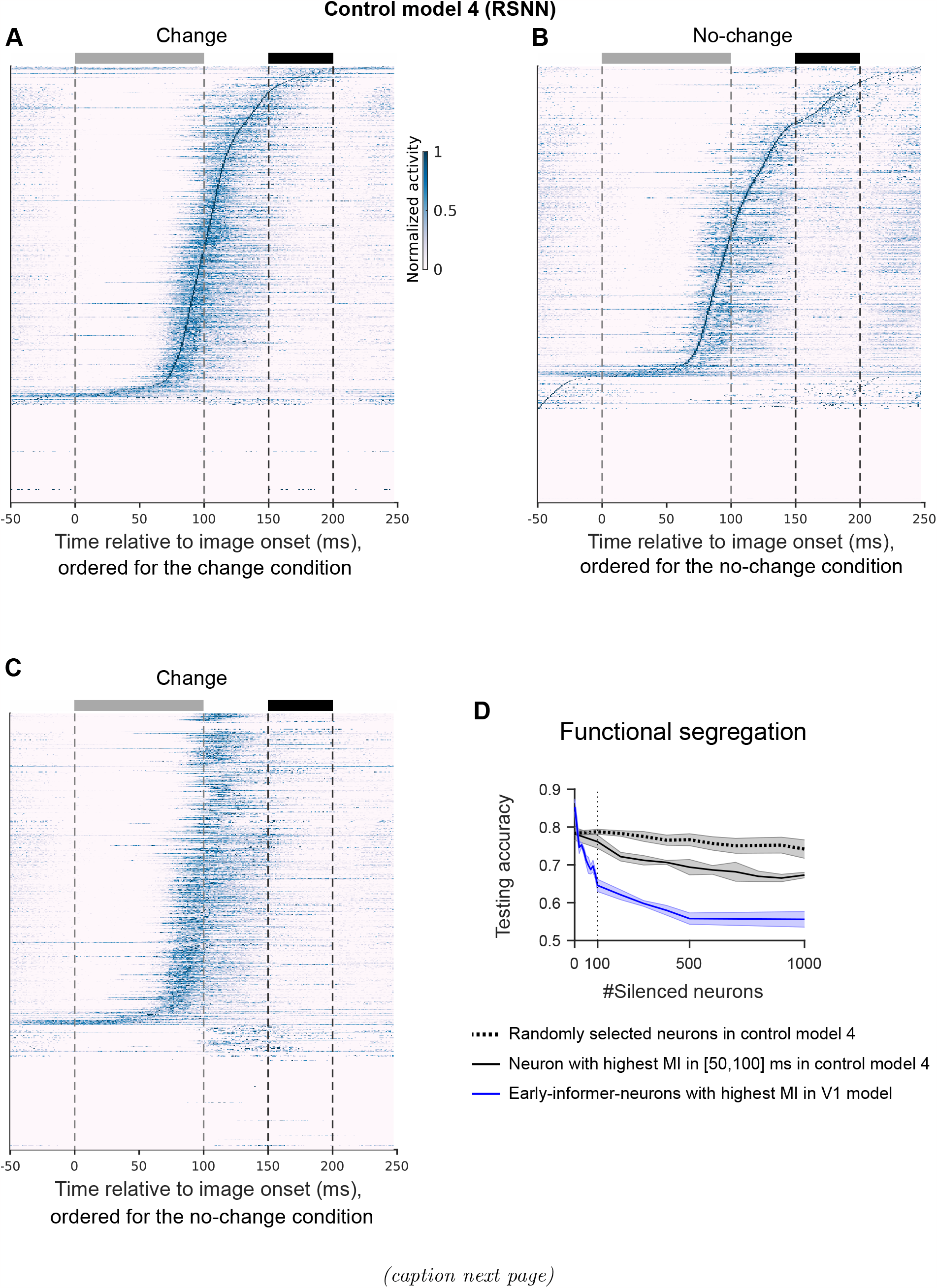
Temporal organization of computations in generic recurrent spiking neural network (RSNN) in visual-change-detection task. **(A)** Normalized average responses of 200 change-condition trials in RSNN are plotted over time, with neurons ordered based on the time of their peak activities under change condition. The gray and black rectangles denote the image presentation and response windows, respectively. In contrast to the V1 model, many neurons are inactive. This is likely due to the need for reducing the loss of firing-rate regularization. **(B)** Same as in **(A)**, but for the no-change condition. **(C)** Normalized neural activities in the change condition are visualized with the order of neurons in the no-change condition, showing large similarity to both **(A)** and **(B)**, unlike the distinction observed in the V1 model. **(D)** Test accuracy as a function of silencing random neurons or early-informer-neurons. All curves show average values for 10 models where different sets of training data were used. The shaded area represents the SEM across 10 models.

**Figure S13:**
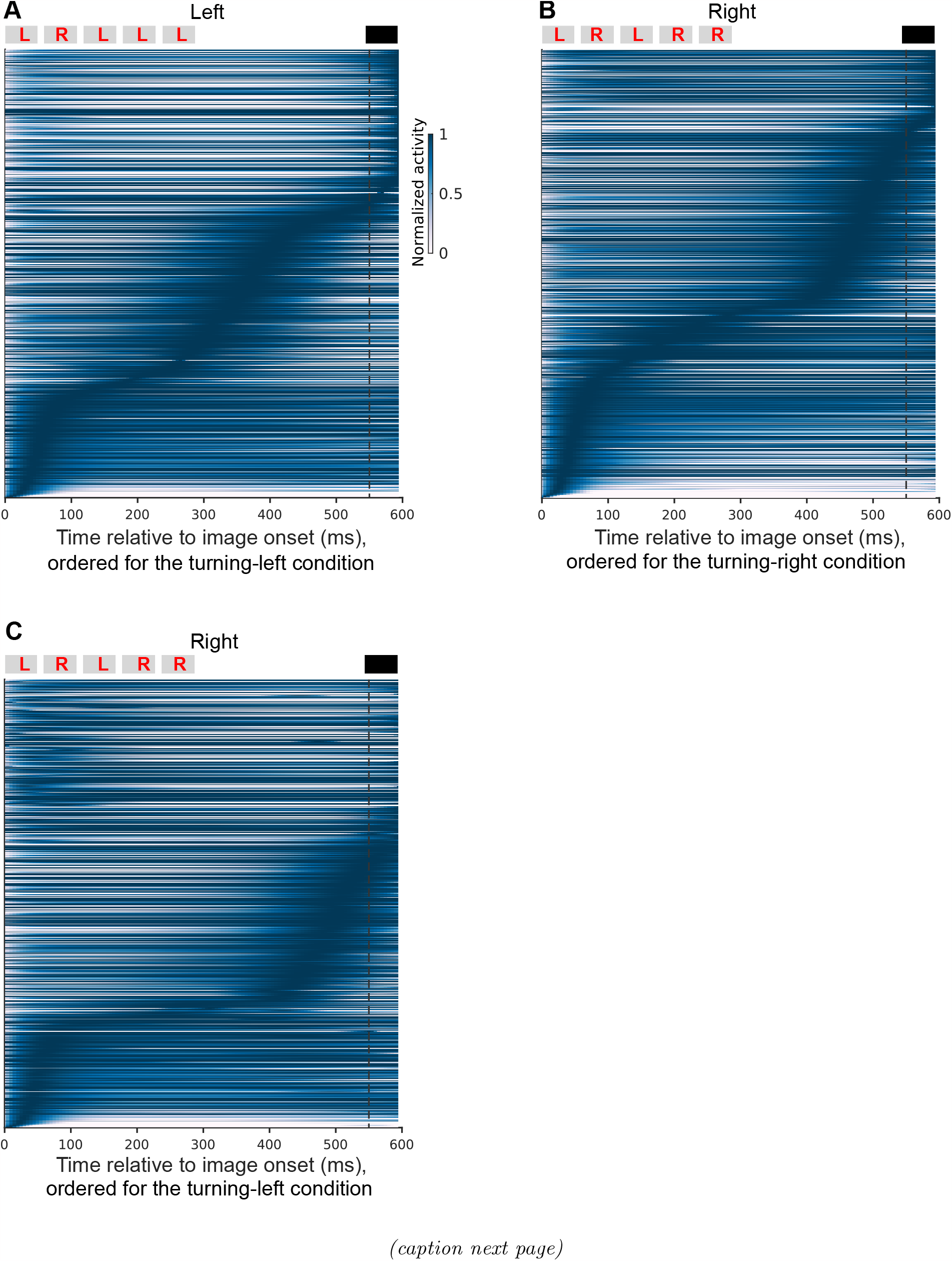
Temporal organization of computations of evidence accumulation task in the RANN model. **(A)** Normalized average responses over 200 trials with the turning-left condition are shown, with neurons ordered according to the time of their peak activity under the turning-left condition. The gray and black bars at the top denote the cue presentations and response windows, respectively. The yellow letters, L (left) / R (right) in gray bars represent the meaning of cue. To reduce the variability, we used the sequence, LRLLL for the turning-left trials. **(B)** Same as in **(A)**, but for the turning-right condition, with neurons ordered according to the time of their peak activity for the turning-right condition. To reduce the variability, we used the sequence, LRLRR for the turning-right trials. **(C)** The same data as in **(B)** for the turning-right condition, but with neurons ordered as in **(A)**. The resulting blurred sequence indicates that the order of peak activity of neurons is quite different for the turning-left and turning-right conditions.

**Figure S14:**
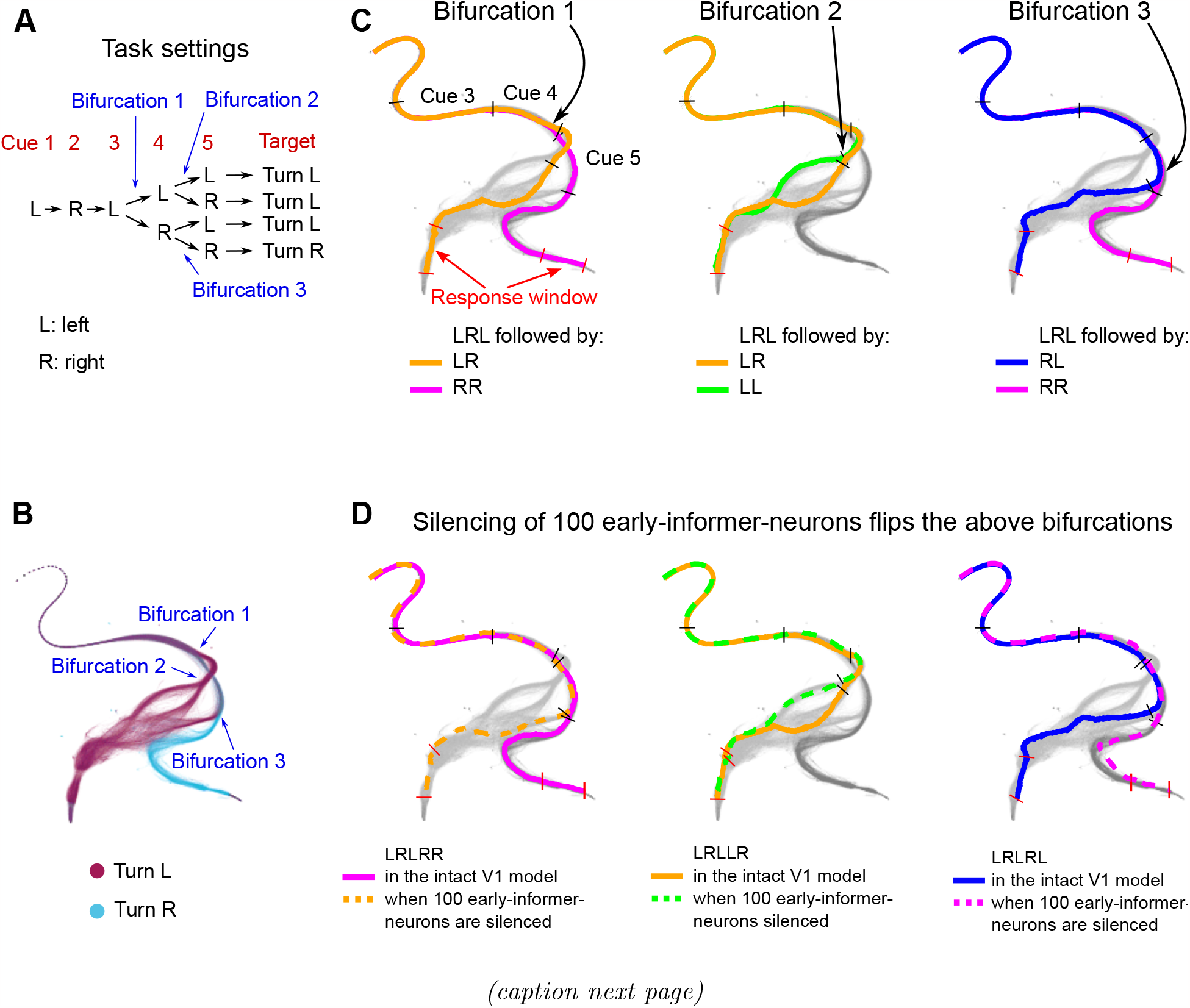
Nested bifurcations of trajectories of network states provide links between network dynamics and network computations in evidence accumulation task. Each dot represents network activity at a particular millisecond of 530-ms (the first 70 ms was excluded for better visualization) fragment of the computation of the V1 model in the evidence accumulation task. Short black bars mark cue 3-5 on the network trajectories; the red bars denote the response window. **(A)** The task setting. The sequence for the first three cues is consistently left-right-left, while the last two cues can be either left or right. The setting introduces three bifurcations. **(B)** Network trajectories of the V1 model during the presentation of five cues. The color differentiation indicates whether the target is to turn left or right. It is evident that the network state trajectories bifurcate three times, depending on the sequence of the fourth and fifth cues. **(C)** The first bifurcation corresponds to the fourth cue, typically occurring 30 ms after its onset. The second bifurcation takes place in response to the fifth cue, provided that the fourth cue is left; however, these two branches eventually converge as they share the same targets. The third bifurcation responds to the fifth cue when the fourth cue is right. **(D)** Those bifurcations in **(B)** can be flipped by a small set of early-informer-neurons. In each sub-panel, there are two trajectories that have the same fourth and fifth cues in both the intact model and when 100 neurons with the highest mutual information with the current condition of accumulated cues in a temporal segment are silenced. For enhanced visualization, the silenced neurons in the panel from left to right are determined based on the mutual information with the current condition of accumulated cues computed during the fourth, fifth, and fifth cues, respectively. These cues align with the bifurcation pivot points. Note that the displayed altered trajectories represent trials in which the network made incorrect decisions.

**Figure S15:**
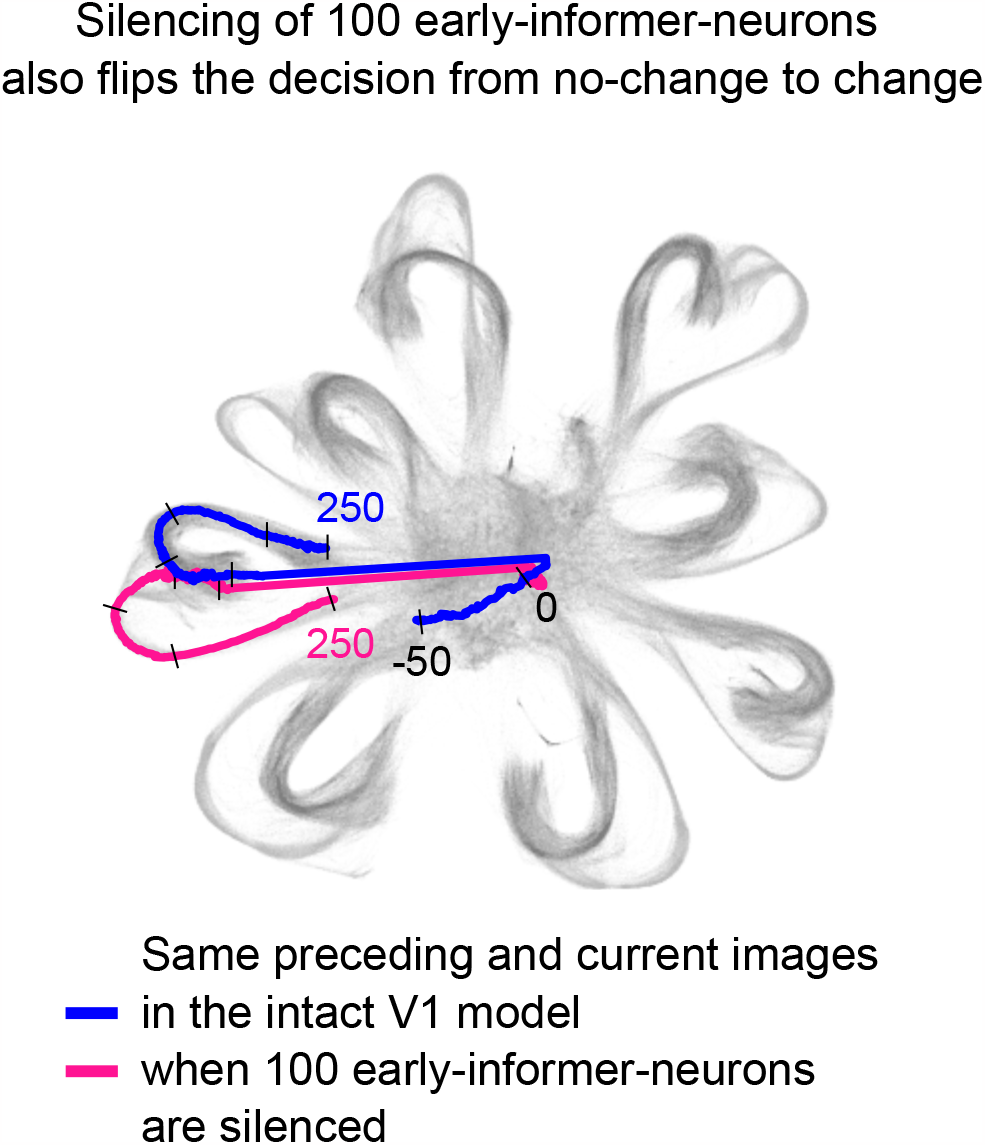
Silencing of 100 neurons can also flip a trajectory from the bundle for no-change to the change bundle of trajectories in the visual-change-detection task. In Fig. 5D, we demonstrated that silencing 100 early-informer-neurons can cause the trajectory of network states to flip from the bundle for change to the bundle for no-change trials. We show here that silencing of the same 100 neurons can also flip the network bifurcation in the other direction.

**Figure S16:**
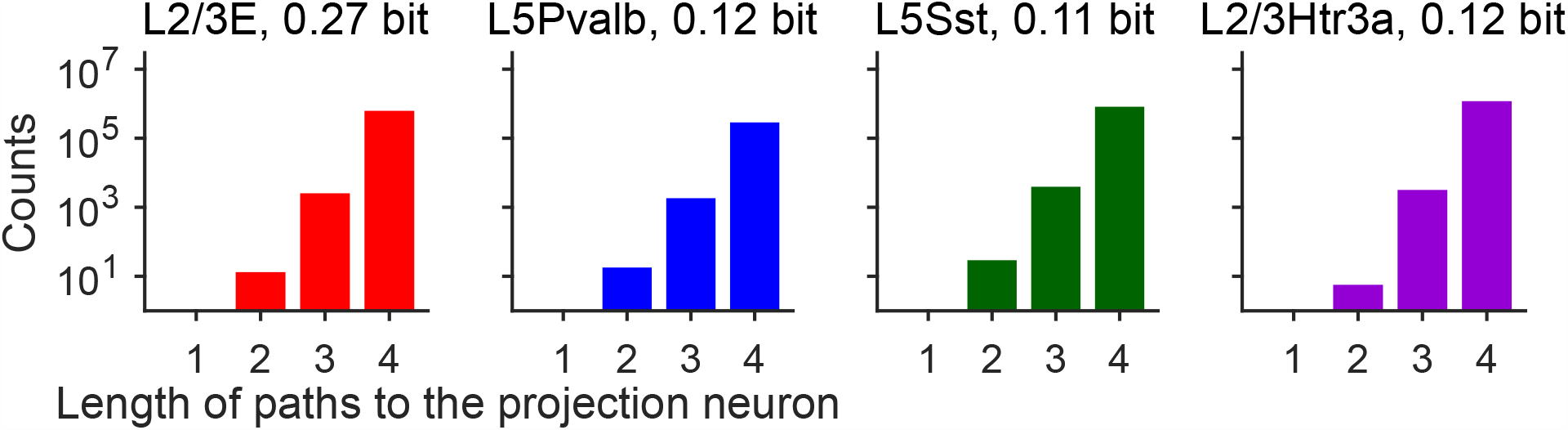
Histogram of the lengths (= number of synapses) of paths from 4 sampled early-informer-neurons with high MI to the projection neuron in the visual-change-detection task. We selected early-informer-neurons with the highest MI in four basic neuron types (excitatory, Pvalb, Sst, and Htra3 neurons). The paths from early-informer-neurons to the projection neuron were found by the MATLAB function “allpaths” in the directed graph. One can also find arbitrarily long paths; here we only demonstrate the short paths (length *<* 5) to the projection neuron for each of the early-informer-neurons. This distribution of path lengths suggests that the firing activity of an early-informer-neuron affects the membrane voltage of the projection neuron in multiple and diverse indirect ways.

